# Structure of the minor spliceosomal U11 snRNP

**DOI:** 10.1101/2023.12.22.573053

**Authors:** Jiangfeng Zhao, Daniel Peter, Irina Brandina, Xiangyang Liu, Wojciech P. Galej

## Abstract

The minor spliceosome catalyses the excision of U12-dependent introns from pre-mRNAs. These introns are rare, but their removal is critical for cell viability. We obtained a cryo-EM reconstruction of the 13-subunit U11 snRNP complex, revealing structures of U11 snRNA and five minor spliceosome-specific factors. U11 snRNP appears strikingly different from the equivalent major spliceosome U1 snRNP. SNRNP25 and SNRNP35 form a dimer, which specifically recognises U11 snRNA. PDCD7 forms extended helices, which bridge SNRNP25 and SNRNP48, located at the distal ends of the particle. SNRNP48 forms multiple interfaces with U11 snRNP and, together with ZMAT5, are positioned near the 5’-end of the U11 snRNA and likely stabilise the binding of the incoming 5’SS. Our structure provides mechanistic insights into U12-dependent intron recognition and the evolution of the splicing machinery.

## Introduction

Precursors of eukaryotic messenger RNAs (pre-mRNAs) contain non-coding segments (introns), which are removed during gene expression by a large and dynamic RNA-protein complex known as the spliceosome(*1*–*3*). The spliceosome assembles *de novo* on each individual intron. A series of complex conformational and compositional rearrangements leads to the formation of the spliceosome RNA catalytic core, allowing it to perform two trans-esterification steps, resulting in spliced mRNA and lariat intron products.

The vast majority of human introns (U2-dependent introns) are processed by the major spliceosome consisting of five canonical subunits, U1, U2, U4, U6 and U5 small nuclear ribonucleoprotein particles (snRNPs) and numerous additional, non-snRNP factors. However, a small class (<1%) of introns (U12-dependent introns) utilise distinct splicing machinery consisting of four unique snRNAs (U11, U12, U4atac and U6atac)(*4*, *5*) and at least 14 unique protein factors(*6*–*10*). These factors work conjointly with the numerous core components shared between the two types of machinery (e.g. U5 snRNP, SF3b complex)(*6*, *7*, *11*) and together assemble into the minor spliceosome. U12-dependent introns are rare but are biased to be located in genes with critical cellular functions, suggesting their ancient origin(*12*, *13*). Consequently, malfunction of the minor spliceosome has severe consequences for human health and is linked to cancer and autoimmune disorders as well as neurodegenerative diseases(*14*).

Although it is believed that the basic mechanistic principles of the minor and major spliceosomes are likely to be conserved(*15*), there are substantial differences between the two systems. U12-dependent introns tend to be shorter, contain no clear polypyrimidine tract and have much stronger splice site sequence conservation compared to the U2-dependent introns (*13*, *16*). During major spliceosome assembly, U1 and U2 snRNP bind sequentially to the pre-mRNA(*17*), forming first an ATP-independent complex E(*18*) followed by an ATP-dependent, stable incorporation of the U2 snRNP into the prespliceosome (complex A)(*19*, *20*). In contrast to that, the U12-dependent intron recognition is carried out by the pre-assembled U11/U12 di-snRNP complex(*21*), which binds cooperatively to both the 5’-splice site (5’-SS) and branch point sequences (BPS)(*22*). U11/U12 di-snRNP contains U11 and U12 snRNAs, two sets of seven Sm proteins, SF3b complex and eight minor spliceosome-specific factors: ZMAT5 (U11-20K), SNRNP25 (U11-25K), ZCRB1 (U11/U12-31K), SNRNP35 (U11-35K), SNRNP48 (U11-48K), PDCD7

(U11-59K), RNPC3 (U12-65K) and ZRSR2 (*6*). Four of these factors show putative sequence homology to major spliceosome proteins, ZMAT5 to U1-C,(*6*), SNRNP35 to U1-70K(*23*), ZRSR2 to U2AF1(*24*) and RNPC3 to SNRPB2/SNRPA(*25*). From the remaining proteins, SNRNP48 has been shown to interact with the U11 5’-SS by site-specific cross-linking and NMR binding studies(*26*, *27*). SNRNP25, SNRNP35, and SNRNP48 co-migrate with the 12S mono U11 snRNP(*6*) and a chain of interactions between SNRNP25-PDCD7-RNPC3 was postulated to bridge the U11 and U12 snRNPs(*25*, *26*).

While the major spliceosome has been extensively studied using biochemistry, genetics and cryo-EM(*28*–*32*), the structural and biochemical information on the minor spliceosome is very scarce. Here, we report a cryo-EM structure of the 13-subunit human minor spliceosome U11 snRNP, revealing the structure of the U11 snRNA and five minor spliceosome-specific proteins (ZMAT5, SNRNP25, SNRNP35, SNRNP48 and PDCD7). Our structure provides mechanistic insights into U12-dependent intron recognition and the evolution of the splicing machinery.

## Results

### The composition and overall architecture of the U11 snRNP complex

We isolated human U11 snRNP from a stable HEK293F cell line ectopically over-expressing ProteinA-tagged SNRNP35, which was used as bait for sample purification and EGFP-tagged ZCRB1, used for sample monitoring (Fig. S1). IgG sepharose affinity purification under stringent, high-salt conditions followed by glycerol gradient allowed us to dissociate the U11/U12 di-snRNP and isolate a stable 13-subunit U11 snRNP complex composed of U11 snRNA, seven Sm proteins, and five minor spliceosome-specific factors ZMAT5, SNRNP25, SNRNP35, SNRNP48, PDCD7 (Table S1). This stable core complex was subjected to single-particle cryo-EM analysis resulting in a 3D reconstruction at the overall 3.4 Å resolution (Figs. S2 and S3; Table S2).

The particle has an elongated, two-lobe architecture consisting of the body of the U11 snRNA bound by SNRNP25 and SNRNP35, and the second lobe containing the Sm ring bound by ZMAT5, SNRNP48 and the 3’-end of the U11 snRNA - stem-loop 4 (SL4) (Fig. 1). The two lobes are connected by extended alpha-helices of the PDCD7, spanning the entire length of the molecule (Fig. 1, Table S3).

**Figure 1.**
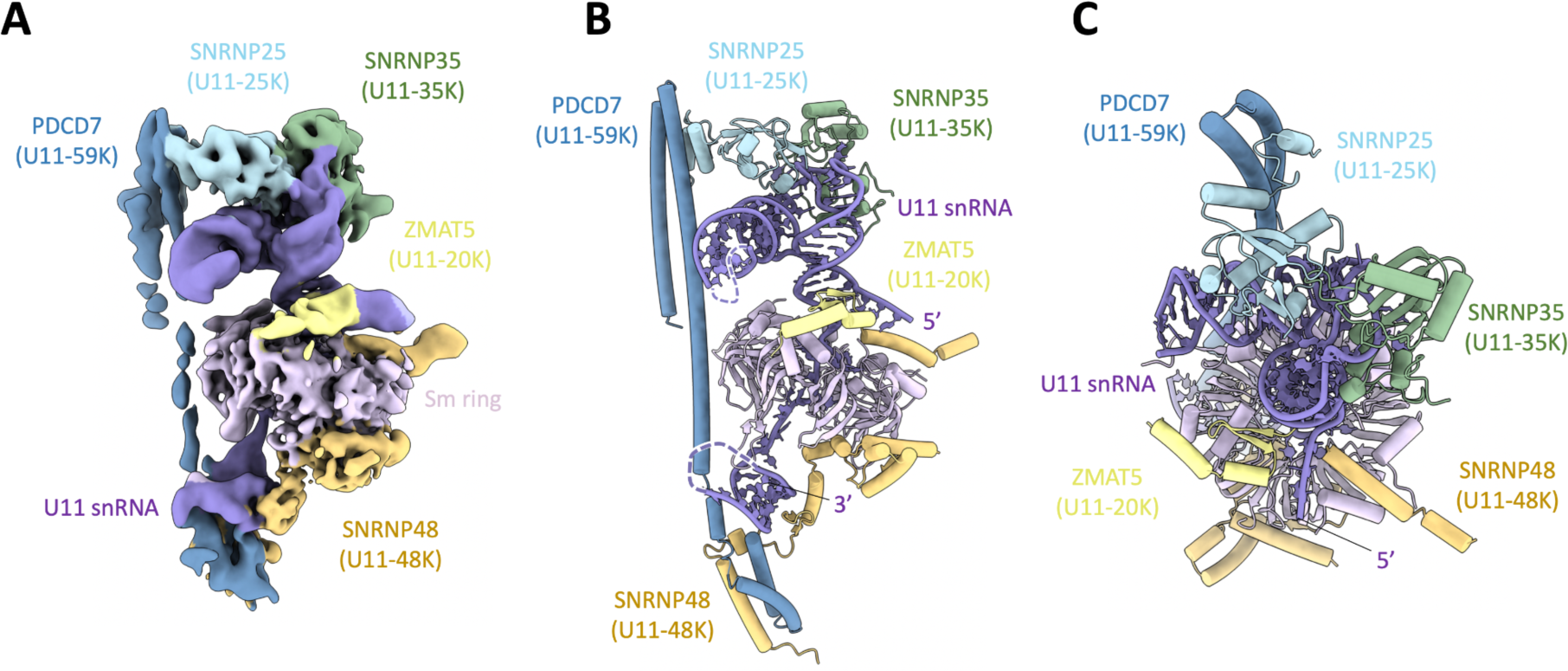
The overall structure of the 13-subunit minor spliceosome U11 snRNP complex. (**A**) Experimental cryo-EM map coloured by the subunit identity. (**B**) Atomic model of the U11 snRNP in the same orientation as in (A). (**C**) Top view of the atomic model.

### Structure of the U11 snRNA

High-resolution cryo-EM reconstruction allowed us to model nearly complete U11 snRNA, including the characteristic 4-helix junction formed between helix H and stem-loops (SL) 1-3 (Figs. 2A-B and S4) as well as the Sm site and SL4, which was modelled at a lower resolution (Figs. S4 and S5). SL1 and SL2 are coaxially stacked and perpendicular to coaxially stacked helices H and SL3 (Fig. 2B). Similar geometry has been observed in yeast and human U1 snRNA(*33*–*35*) (Figs. 2C and S6). One striking feature of the U11 snRNA is the stacking interaction between the bases located in the loops of SL1 and SL3, bringing both stem-loops into close proximity (Figs. 2D-F and S4). Corresponding helices of the U1 snRNP are too long to form a similar arrangement. This unique interaction between U11^SL1^ and U11^SL3^ forms the structural basis for the specific recruitment of minor spliceosome proteins. The 5’-SS binding region of the U11 snRNA is not well resolved in the cryo-EM map, however, a continuous density can be traced from the bottom of the helix H pointing towards the outside of the Sm ring (Fig. 1). This is in agreement with the equivalent region in the U1 snRNP structure(*33*, *36*, *37*). The U11^SL4^ was largely modelled by rigid-body docking and modifying the corresponding parts of the U1 snRNP(*36*), as the map of this region is too noisy for accurate *de novo* modelling.

**Figure 2.**
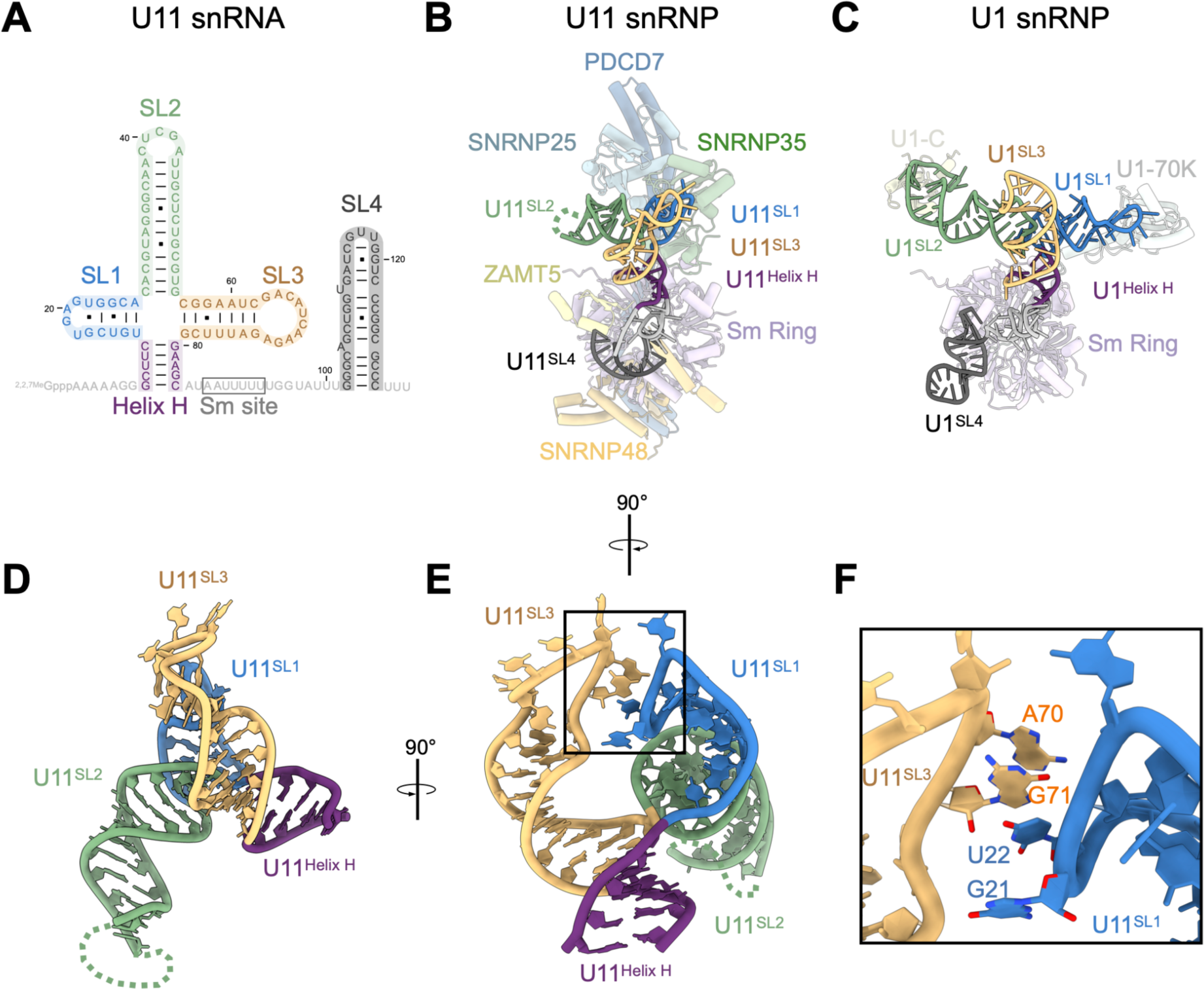
Structure of the U11 snRNA. **(A)** Secondary structure model of the human U11 snRNA, based on the atomic coordinates. **(B)** U11 snRNP structure with the U11 snRNA highlighted in the foreground. Secondary structure elements (Stem-loops) are coloured as in panel (A). **(C)** The structure of the human U1 snRNP (PDB ID: 7B0Y(*55*)) is shown in the same orientation as the U11 snRNP in panel (B) with U1 snRNA highlighted in the foreground. **(D-E)** Two orthogonal views of the atomic model of the U11 snRNA. **(F)** Zoom in on the stacking interaction between the bases in the SL1 and SL3.

### Specific recognition of the U11 snRNA

Two core components of the U11 snRNP, SNRNP35 and SNRNP25, form a heterodimer via a loop extending from the canonical SNRNP35^RRM^ domain. Together, both proteins specifically recognise the unique arrangement of U11^SL1^-U11^SL3^ (Figs. 3A-D and S5). Our structure reveals that SNRNP25 contains a Ubiquitin-like domain (SNRNP25^UBL^) and an N-terminal extension responsible for PDCD7 binding, referred to as the PDCD7-binding domain (SNRNP25^PBD^) (Fig. 3E). The SNRNP25^UBL^ domain overlays very well with the Ubiquitin fold (RMSD=1.5Å with PDB:1UBQ), except for an extension of one alpha-helix, which was adapted to bind RNA at the U11^SL1^ and U11^SL3^ interface (Fig. 3F).

**Figure 3.**
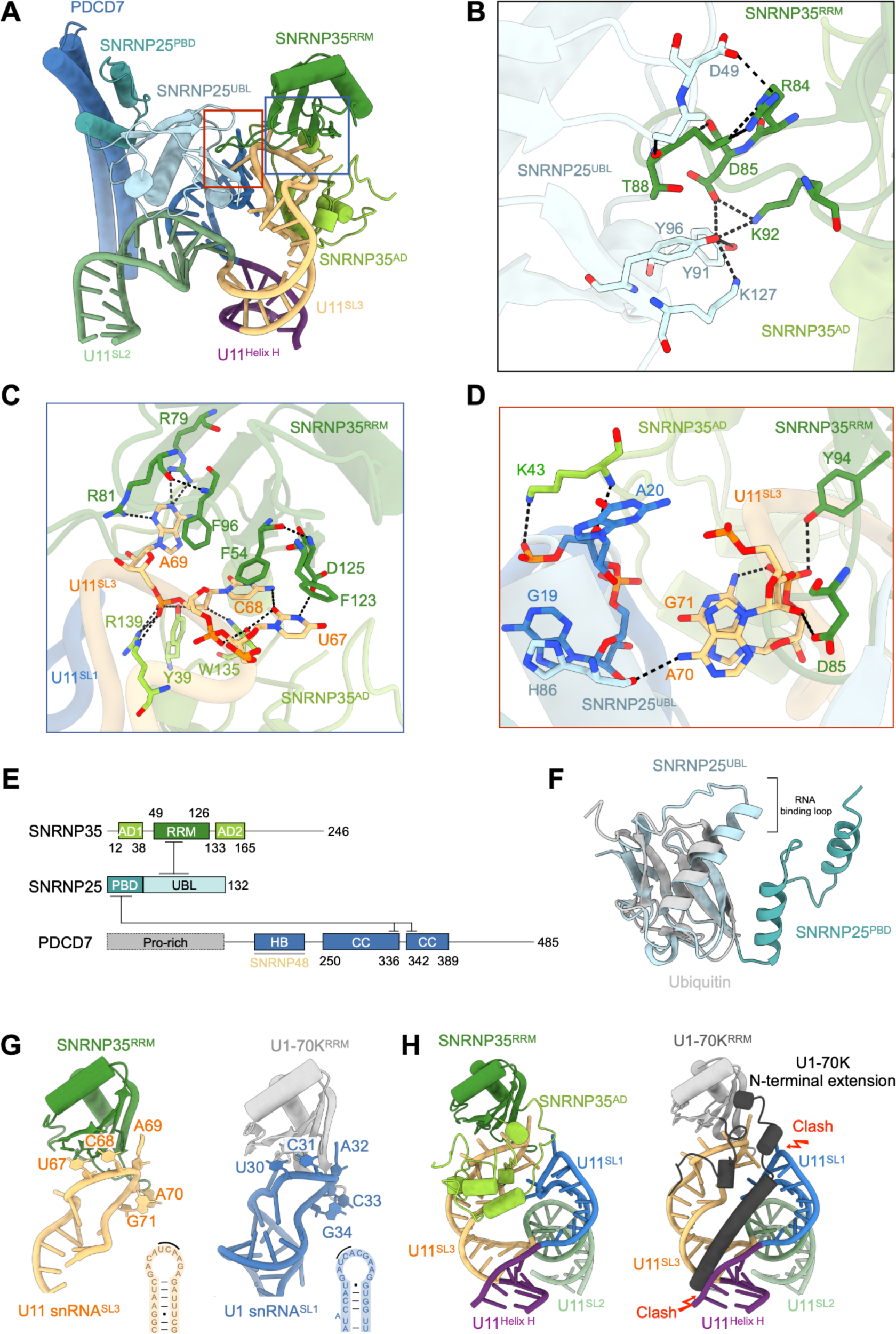
Specific recognition of the U11 snRNA by SNRNP25 and SNRNP35. (**A**) Overall architecture of the U11 snRNA binding region. (**B**) Close view of the interaction between SNRNP25 and SNRNP35. The loop extruded from SNRNP35^RRM^ forms several hydrogen bonds with SNRNP25^UBL^ domain. (**C**, **D**) Zoom in on the recognition of the unique arrangement of U11^SL1^-U11^SL3^ loop region. (**E**) Domain architectures of SNRNP35, SNRNP25 and PDCD7; AD1 - accessory domain 1; AD2 - accessory domain 2; RRM - RNA recognition motif; PBD - PDCD7-binding domain; UBL - Ubiquitin-like domain; HB - Helix bundle; CC - coiled coil. Regions of each protein interacting with each other are indicated. (**F**) Comparison between SNRNP25^UBL^ domain with typical Ubiquitin fold (PDB: 1UBQ). (**G**) Canonical RNA-RRM binding mode of SNRNP35:U11^SL3^ and U1-70K:U1^SL1^. The secondary structures of corresponding stem loops are shown in the right-bottom corner, and the conserved 5’-UCA-3’ sequence is indicated by black curve. (**H**) Specific recognition of U11 snRNA by SNRNP35, rather than U1-70K. The N-terminal helix of U1-70K would clash with the U11^SL1^.

U1-70K, which was previously identified as a homologue of the SNRNP35(*23*), uses its RRM domain to specifically bind the SL1 of the U1 snRNA (U1^SL1^)(*36*, *38*). In the U11 snRNP, the SL1 is much shorter and cannot form similar interactions. SNRNP35^RRM^ domain binds to U11^SL3^ instead. Interestingly, despite binding to different stem-loops, SNRNP35^RRM^ and U1-70K^RRM^ domains form similar stacking interactions with the conserved 5’-UCA-3’ sequence splayed on top of the RRM’s RNP motifs, in agreement with the canonical RNA-RRM binding mode(*39*, *40*)(Fig. 3G). Much of the RNA binding specificity of the U1-70K is achieved thanks to large extensions flanking the RRM domain(*36*). Similarly, SNRNP35 contains extensions on both sides of the canonical RRM domain, that form a composite accessory domain (SNRNP35^AD^) binding specifically to the base of the U11^SL3^ and apical region of the U11^SL1^ (Fig. 3C, D). Importantly, the extensions of the SNRNP35 and U1-70K RRM domains confer the specificity for the integration of each protein in their cognate splicing systems (i.e. U1-70K cannot bind to U11 snRNA as it would clash with the U11^SL1^) (Fig. 3H).

### PDCD7 mediates long-range interactions between distal ends of the U11 snRNP

One of the most striking features of the U11 snRNP structure are the extended coiled coils of the PDCD7 (PDCD7^CC^), which are tethered to the U11 snRNA via the SNRNP25^PBD^ domain. These long alpha helices run along the entire length of the particle (over 170Å), reaching the Sm ring and U11^SL4^ (Fig. 1). The N-terminal extension of the tip of the coiled-coil region of PDCD7 (residues 171-225) forms a three-helix bundle together with the SNRNP48 (residues 8-32) and is located at the bottom of U11^SL4^ (Fig. S4). Although the resolution of our map is limited in this region, such an arrangement is consistent with AlphaFold2 structure prediction(*41*, *42*) and yeast two-hybrid experiments showing the interaction of the PDCD7(residues 137-336) with SNRNP48(*26*). Based on this observation, PDCD7 likely recruits the N-terminus of SNRNP48 near the U11^SL4^ and facilitates its binding to other components of the U11 snRNP. Most notably SNRNP48 zinc finger domain (SNRNP48^ZnF^) binds to the minor groove of the U11^SL4^, the Sm binding domain (SNRNP48^SBD^) to the side of the Sm ring, and the C-terminal helix (SNRNP48^Cterm^) interacts with SmE, near the 5’-end of U11 snRNA (Figs. 4A and S4). Since the interactions between SNRNP48 and the Sm ring are, in principle, compatible with any other Sm-containing snRNPs, it appears that PDCD7-mediated recruitment of SNRNP48 confers the specificity for its association with U11 snRNP.

**Figure 4.**
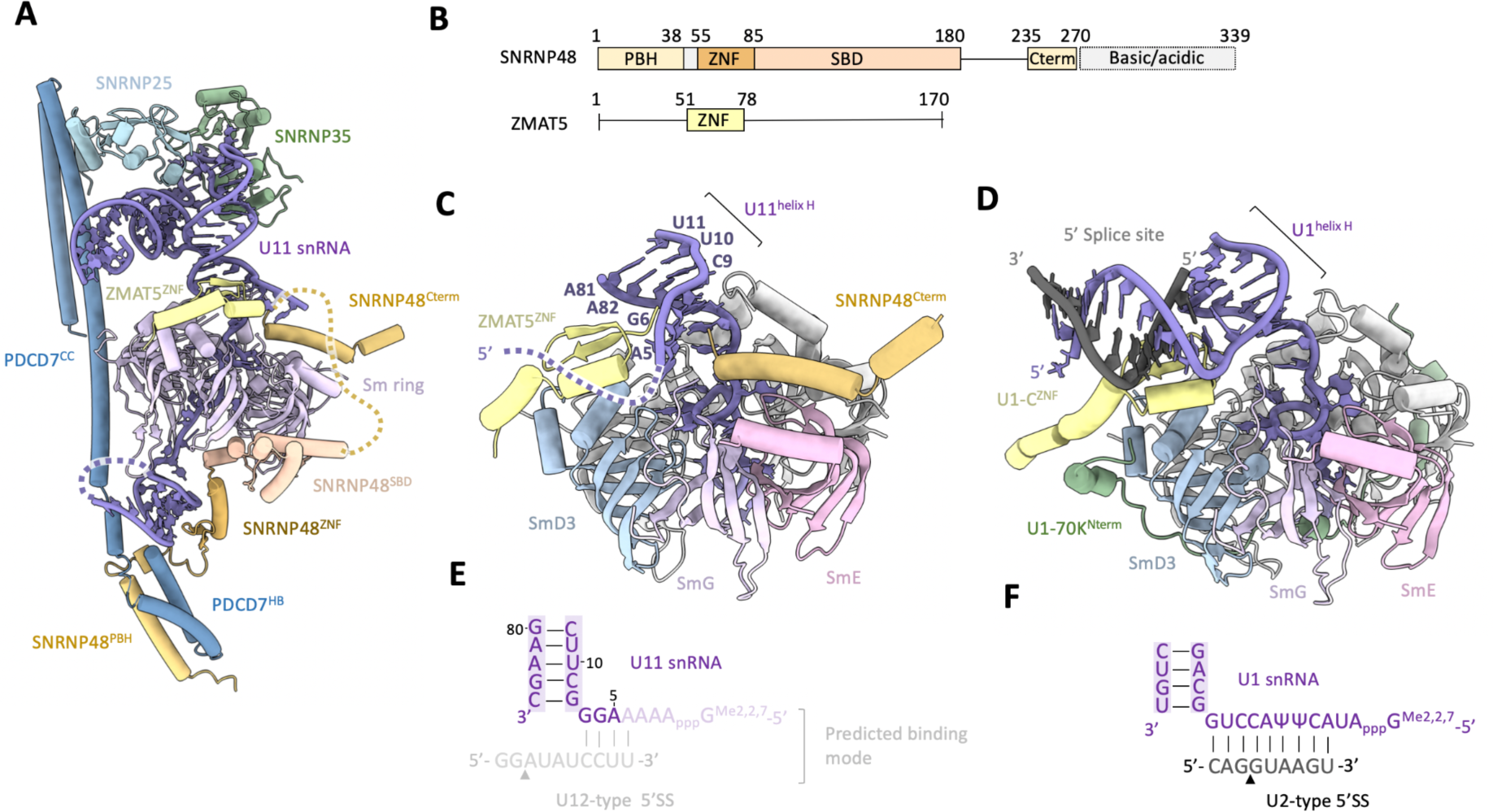
SNRNP48 recruitment by PDCD7 and its implications for the 5’SS recognition. **(A)** SNRNP48 forms multiple contacts with the U11 snRNP components. **(B)** Domain architectures of ZMAT5 and SNRNP48; PBH - PDCD7-Binding Helix; CC - coiled coil; ZNF - Zinc Finger domain; SBD - Sm Binding Domain. **(C)** 5’SS binding region of the U11 snRNA is positioned between ZMAT5^ZNF^ and SNRNP48^Cterm^ domains, suggesting their possible involvement in the stabilisation of the 5’SS binding. **(D)** an equivalent view of the U1 snRNP bound to the model 5’SS(*36*). **(E)** The secondary structure of the RNA shown in (C). A putative 5’SS binding mode was included (lighter colours) for the sake of the comparison with U1, however, it was not resolved in the current structure. **(F)** The secondary structure of the RNA shown in (D).

### Implications for the U12-type 5’SS recognition

The 5’SS binding region of the U11 snRNA extends from the helix H towards the periphery of the Sm ring (Figs. 1 and 4). It is surrounded by two tubular densities, which were interpreted as the C-terminus of SNRNP48 and the Zinc finger domain of ZMAT5 (ZMAT5^ZNF^) (Figs. S7 and S8). ZMAT5 has been previously reported to show some homology to the U1-C(*6*), and its interaction with SmD3 and location directly next to 5’SS binding region of the U11 snRNA indeed resembles the arrangement observed for the U1 snRNP(*33*) (Figs. 4 and S4). U1-C interacts with the phosphate backbone of the U1 snRNA:5’SS duplex and increases their affinity in a sequence-independent manner(*36*). It is likely that ZMAT5^ZNF^ plays a similar role in the minor 5’SS recognition.

On the other side of the 5’SS binding region, the C-terminal helix of the SNRNP48 is positioned near the 5’-end of U11 snRNA via its interface with the SmE. SNRNP48 has been shown to interact with the 5’SS (U+2) by site-specific cross-linking(*26*). This observation is compatible with SNRNP48^Cterm^ location in our structure, but the details of such an interaction remain to be investigated in the substrate-bound U11 snRNP. Interestingly, part of the SmE interface used to recruit SNRNP48^Cterm^ overlaps with the position of the N-terminal helix of Luc7, as visualised in the yeast pre-spliceosome(*37*) (Fig. S9). While both proteins share the same binding site, their architectures are very different, and it is unlikely that they would act in a similar fashion (Fig. S9).

### Bridging interaction between U11 and U12 snRNPs

U11/U12 di-snRNP exists in cells as a stable dimer(*21*). While stringent purification conditions allowed us to isolate U11 snRNP core complex and solve its structure, the information concerning U11-U12 interactions has been partially lost. Nevertheless, we noted that RNPC3 is present in our sample and it was shown in yeast two-hybrid experiments to be an interactor of the PDCD7(residues 332-481)(*25*). Since RNPC3 also binds the U12 snRNA, it was postulated to act as a bridge between the U11 and U12 snRNPs (*25*, *43*). AlphaFold2 predicts a high-confidence interface between PDCD7(residues 415-450) and a disordered region of RNPC3 (residues 175-190) as well as the N-terminus of ZMAT5 (Fig. S10). Our cryo-EM reconstruction shows fuzzy density around the C-terminal α-helix of the PDCD7^CC^ domain. This density could not be interpreted unambiguously, but it is consistent with a putative position of the predicted RNPC3:PDCD7:ZMAT5 composite domain (Fig.5 and S10). While we could map the RNPC3 binding site on U11 snRNP, the exact location of its RRMs and putative position of U12 snRNP with respect to U11 snRNP cannot be reliably modelled due to extended linkers between RNPC3^RRM^ domain and the PDCD7 binding site. It is possible that such a flexible arrangement could be functionally important for binding to a wide range of pre-mRNA substrates.

**Figure 5.**
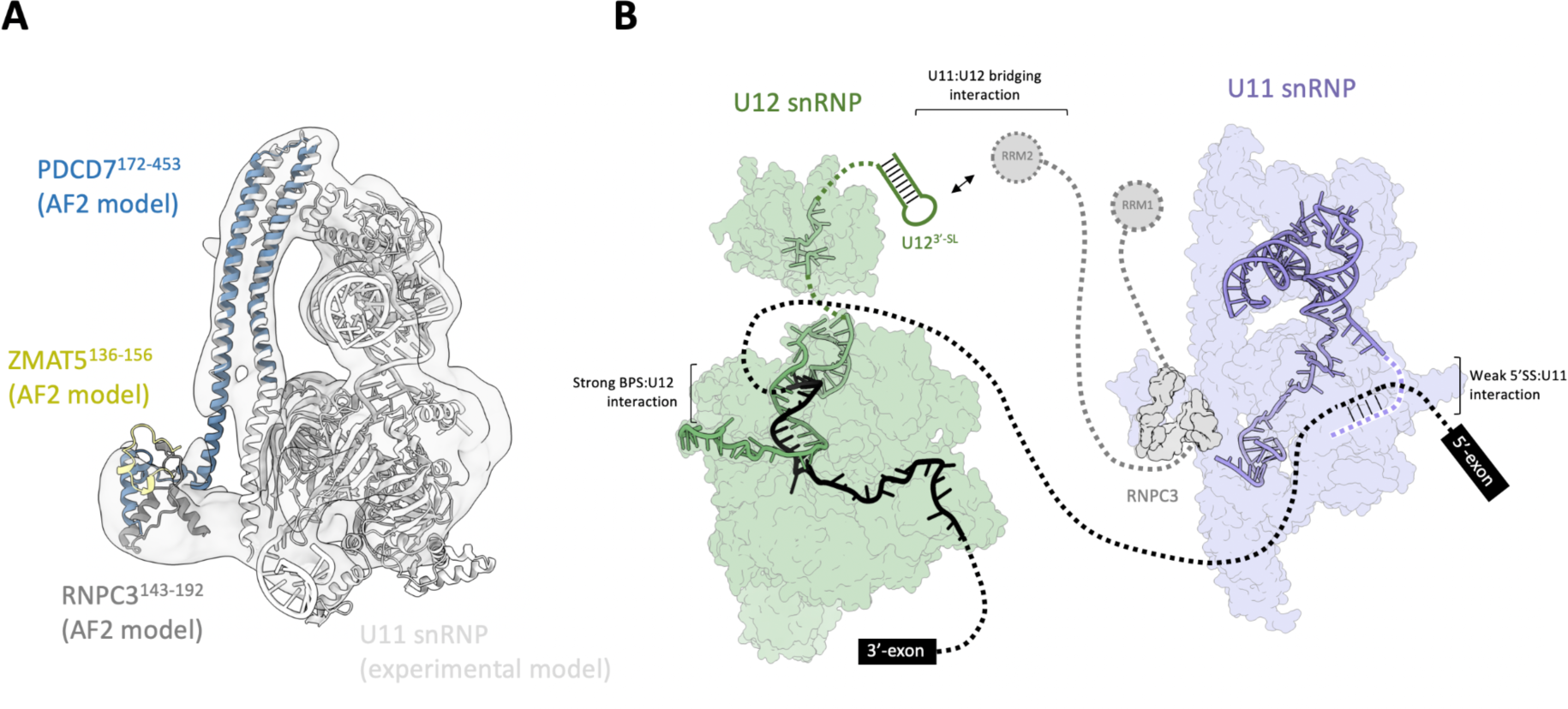
Bridging interactions between U11 and U12 snRNPs. **(A)** AlphaFold2 (AF2) prediction of the ternary PDCD7:ZMAT5:RNPC3 complex, coincides with the low-resolution unassigned density when superimposed via PDCD7 coiled-coil domain. **(B)** A model of the RNPC3-mediated U11 snRNP recruitment to the U12 snRNP.

## Discussion

U12-dependent introns are widely spread across eukaryotes, however, the mechanism of their recognition and removal by the minor spliceosome remains elusive. Our structure of the 13-subunit U11 snRNP provides insights into the assembly principles and architecture of the complex containing five minor spliceosome-specific factors, highlighting unique features of the U12-dependent splicing system. U11 snRNP is strikingly different from the U1 snRNP, which plays an equivalent role in the U2-dependent spliceosome. With the exception of the Sm ring (shared with the U1) and ZMAT5^ZNF^, which shows a clear resemblance to U1-C, none of the other components have counterparts in the major splicing system. This concerns even SNRNP35, previously postulated to be related to the U1-70K, which shows a very different binding topology despite the conservation of the canonical RNA-RRM binding mode.

These unique features are most likely a manifestation of the need for specific recognition of the minor spliceosome components in the presence of much more abundant major splicing factors. This is best exemplified by the SNRNP25-SNRNP35 dimer, which forms several complex interfaces with the RNA elements unique to the U11 snRNA. Importantly, this specific recognition is transmitted to distal parts of the complex via a chain of interactions starting with SNRNP25, PDCD7 through SNRNP48 and ZMAT5 all the way to the proximity of the 5’SS binding region. There is a precedence for such a long-distance interaction in the U1 snRNP, where an extended tail of the U1-70K wraps around the Sm ring and contacts U1-C near the 5’SS(*33*, *44*). However, the components and the mechanism utilised by the U11 snRNP to achieve similar goals are very different. Notably, the unique architecture of the U11 snRNA is incompatible with the binding of U1-specific proteins, preventing potential cross-reactivity of the components of both splicing systems.

5’ splice sites of the U12-type introns have a highly conserved consensus sequence, but their base-pairing potential to U11 snRNP is limited. Our structure shows that only four nucleotides of the previously identified 5’SS binding region of the U11 snRNA(*26*, *45*, *46*) remain single-stranded and available for base-pairing. This contrasts with a much longer 5’-SS binding region of the U1 snRNP (*47*). Conservation of the 5’SS region not involved in the U11 snRNP binding could be explained by its interaction with U6atac and between the 5’SS and 3’SS during catalytic stages of splicing(*7*, *48*, *49*). It is likely that the two proteins located in the proximity of the U11 5’-end, ZMAT5 and SNRNP48, act in concert to stabilise U11:5’SS interaction, much like it is done by the U1C in the human U1 snRNP(*36*). U1C stabilises 5’SS binding in a sequence-independent manner, but it is not clear if this is also the case for ZMAT5 and SNRNP48, or if they additionally contribute to the specificity of the 5’SS recognition. We noted that part of the binding site of SNRNP48^Cterm^ on the Sm ring is occupied by a conserved α-helix of Luc7 in yeast pre-spliceosome, where it stabilizes the 5’-SS:U1 snRNP interaction via its two zinc finger domains(*37*, *50*, *51*). In humans, three homologs of Luc7, (LUC7L1, LUC7L2, LUC7L3), are involved in alternative splicing regulation(*52*, *53*), differentially affecting distinct sub-classes of the 5’-SS(*54*). The architecture of SNRNP48 is very different from the LUC7L(1-3), and it is unlikely that it would use a similar mechanism to stabilise U11:5’SS interaction (Fig. S9). Nevertheless, as a stable component of U11 snRNP, the SNRNP48^Cterm^ domain may act as a placeholder to prevent unwanted or pre-mature, LUC7L(1-3)-mediated alternative splicing regulation of the U12-type 5’ splice sites.

A very limited base-paring potential of the U11 snRNA and stable association of U11/U12 di-snRNP suggest that the main driving force for the minor intron recognition likely relies on the U12:BS interaction which brings the U11 snRNP into the proximity of the 5’SS and facilitate their interaction (Fig. 5). Such a scenario would help rationalize the evolutionary pressure to maintain U11 and U12 stably associated with each other and their cooperative binding to the target introns(*22*).

## Acknowledgments

We would like to acknowledge Sarah Schneider, Romain Linares, Joseph Bartho, Felix Weis, Wim Hagen and Zhengyi Zhang for their long-term support at the EMBL Electron Microscopy Facilities in Grenoble and Heidelberg; Mandy Rettel, Per Haberkant, and Frank Stein from the EMBL Proteomic Core Facility for the mass spectrometry data acquisition and analysis; ESRF for the provision of the beamtime at CM01 and Isai Kandiah for the assistance with the preliminary data collection; Angelique Fraudeau and Estelle Marchal for the laboratory technical assistance; Martin Pelosse and Alice Aubert from the EMBL Grenoble EEF platform for the assistance with cell culture; A. Peuch and the EMBL Grenoble IT team for the support with high-performance computing; Ramesh Pillai for the critical comments on the manuscript.

## Funding

European Molecular Biology Laboratory (WPG)

European Research Council (ERC) under the European Union’s Horizon 2020 research and innovation programme, grant agreement no. 950278 (WPG)

Human Frontiers Science Program, HSFP Postdoctoral Fellowship LT000383/2018-L (DP)

## Author contributions

Conceptualization: JZ, DP, WPG

Methodology: JZ, DP, IB, XL

Investigation: JZ, DP, IB, XL, WPG

Visualization: JZ, WPG

Funding acquisition: WPG,

Project administration: JZ, WPG

Supervision: WPG

Writing – original draft: JZ, WPG

Writing – review & editing: JZ, DP, IB, XL, WPG

## Competing interests

Authors declare that they have no competing interests

## Materials and Methods

### Generation of a stable cell line overexpressing proteinA-tagged SNRNP35 and EGFP-tagged ZRCB1

Open reading frame of SNRNP35 was cloned into a modified pFLAG_CMV10 vector containing an N-terminal ProteinA-TEV-3×HA affinity tag. The open reading frame of ZCRB1 was cloned into a modified pFLAG_CMV10 vector containing an N-terminal 3×HA-EGFP-3C-SBP affinity tag. Freestyle 293-F cells were co-transfected with these two plasmids, and a stable, polyclonal cell line was derived through Geneticin and Puromycin antibiotic selection. Expression of the target proteins was confirmed by western blot analysis.

### Purification of the U11 snRNP

Modified Freestyle 293-F cell line was grown in the Freestyle medium to the density 2 × 10^6^ cells/ml, and the nuclear extract was prepared following the original Dignam protocol(*56*, *57*). For each preparation, an aliquot of flash-frozen nuclear extract was thawed on ice, and salt concentration was adjusted to the final 500 mM KCl. IgG Sepharose 6 Fast Flow affinity resin (Cytiva) was added to 5% (v/v) of the reaction volume and incubated overnight at 4°C on a turning wheel. The resin was washed with 20-column volumes (CV) of the purification buffer (20 mM HEPES-KOH, pH 7.9, 500 mM KCl, 2 mM MgCl_2_, 0.1% NP-40, 5% (v/v) glycerol) and eluted by adding purification buffer supplemented with 14.5% (v/v) TEV protease and incubating for 2 hours at room temperature.

The eluate was loaded onto 4 ml 10-30% (v/v) glycerol gradients containing 20 mM HEPES-KOH, pH 7.9, 200 mM KCl, 2 mM MgCl_2_. For cryo-EM studies, gradients also contained 0 – 0.05% glutaraldehyde gradient, as in the Grafix protocol(*58*). Gradients were prepared using the Biocomp 108 Gradient Mixer (Biocomp). After 15 hours of centrifugation at 126,000 *×g*, gradients were fractionated into 27 fractions of 150 µl and the cross-linker was quenched with 10 mM Tris-HCl, pH 7.5. Fractions containing the U11 snRNP were identified by SDS-PAGE using the Novex Tris-Glycine system (Thermo Fisher Scientific). The sample was concentrated using 100 kDa MWCO Amicon concentrators (Merk KGaA, Darmstadt) and then dialysed for 1h at 4°C against buffer containing 20 mM HEPES-KOH, pH 7.9, 200 mM KCl, 2 mM MgCl_2_ and used directly for grid preparation without further manipulations.

### Protein identification via LC-MS/MS

Complexes were purified as described using glycerol gradients without crosslinker, and 50 µl of peak fractions were prepared for LC-MS/MS using the SP3 protocol(*59*). All reagents for LC-MS/MS were prepared in 50 mM HEPES pH 8.5. First, cysteines were reduced using 10 mM dithiothreitol at 56°C for 30 minutes. Samples were kept at 24°C and alkylated with 20 mM 2-chloroacetamide at room temperature in the dark for 30 minutes. They were digested with trypsin (Promega), and the peptides were cleaned up using OASIS HLB μElution Plate (Waters). The outlet of an UltiMate 3000 RSLC nano-LC system (Dionex) fitted with a trapping cartridge (μ-Precolumn C18 PepMap 100, 5μm, 300 μm i.d. × 5 mm, 100 Å) and an analytical column (nanoEaseTM M/Z HSS T3 column 75 μm × 250 mm C18, 1.8 μm, 100 Å, Waters) was coupled directly to the Orbitrap Fusion Lumos (Thermo Fisher Scientific) mass spectrometer using the nanoFlex source. The mass spectrometer was operated in positive mode with the capillary temperature set at 275°C. The peptides were introduced into the mass spectrometer via a Pico-Tip Emitter (360 μm OD × 20 μm ID, 10 μm tip) with an applied spray voltage of 2.4 kV.

With the Orbitrap mass spectrometer in profile mode, in the 300-1500 m/z mass range, full mass scans were acquired with a resolution of 120000. The filling time was set to a maximum of 250 ms with a limit of 2e5 ions. The instrument was operated in data-dependent acquisition (DDA) mode, and MSMS scans were acquired in the Iontrap with the scan rate set to rapid and normal mass range, with a fill time of up to 35 ms. A normalised collision energy of 30 was applied. MS2 data was acquired in centroid mode.

To analyse the data, IsobarQuant5(*60*) and Mascot v2.2.07 (Matrix Science) were used. Data were searched against the UniProt Homo sapiens proteome database (UP000005640), which also contained common contaminants and reversed sequences. The following modifications were included in the search parameters: Carbamidomethylation (C) (fixed modification), Acetylation (Protein N-term) and Oxidation (M) (variable modifications). For the full scan (MS1) a mass error tolerance of 10 ppm, and for MS/MS (MS2), spectra of 0.02 Da were set. Trypsin was set as a protease with a maximum of two missed cleavages, the minimum peptide length was seven amino acids, and at least two unique peptides were required for protein identification. The false discovery rate on peptide and protein levels was set to 0.01.

### Negative-staining EM

Negative-stain EM was used to check fractions of the GraFix gradient. For the preparation of negative-stain grids, CF300-Cu (Electron Microscopy Sciences) were glow-discharged for 90 s at 30 s ccm, 100% power with a mixture of 90% argon and 10% oxygen using the Fischione 1070 Nanoclean plasma cleaner. 3.5 μl of each sample was applied and incubated for 60 s, then washed twice in a drop of water and incubated for 60 s in uranyl acetate solution. All liquid was blotted away, and the grids were air-dried. The grids were imaged using a Tecnai G2 Spirit BT microscope (Thermo Fisher Scientific) operated at 120 kV.

### Cryo-EM sample preparation

Concentrated and dialysed Grafix gradient fractions were used for cryo-EM analysis. UltrAufoil Au 200 mesh R2/2 grids (Quantifoil) EM grids were glow-discharged on each side for 90 s at 30 sccm, with 100% power with a mixture of 90% argon and 10% oxygen using the Fischione 1070 Nanoclean plasma cleaner. 3.5 μl of the sample was applied to glow-discharged grids, blotted for 5 s at -7 blot force, 4°C, 90% humidity and plunge-frozen into liquid ethane using Vitrobot Mark IV (Thermo Fisher Scientific).

### Cryo-EM data collection and analysis

Grids were initially screened using a Glacios Cryo-TEM equipped with a Falcon 4i Direct Electron Detector and SelectrisX energy filter (Thermo Fisher Scientific). Final data collection was performed on a Titan Krios TEM (Thermo Fisher Scientific) operated at 300 kV, equipped with a SelectrisX energy filter (Thermo Fisher Scientific) and a Falcon 4i Direct Electron Detector (Thermo Fisher Scientific) at the EMBL Heidelberg cryo-EM service platform(*61*). A magnification of 165,000× was used, corresponding to a pixel size of 0.73 Å/pixel. Automated data acquisition was performed using SerialEM(*62*). 13,809 movies (dataset 1) were recorded with 41.73 e/Å^2^ total dose, with defocus values from -0.7 to -1.7 μm (0.1 μm steps). Another 17,992 movies (dataset 2) were recorded with 42.61 e/Å^2^ total dose, with defocus values from -0.7 to -1.6 μm (0.1 μm steps).

All image processing was performed within cryoSPARC v4.3.1(*63*). Both datasets were motion corrected using cryoSPARC’s implementation of Patch Motion Correction(*63*) with default settings followed by CTF estimation with Patch CTF. Particles were picked by Topaz(*64*) using a custom-trained model. 7,144,222 particles were extracted with four-fold binning (440 pixel original box size, 110 pixel binned box size, 2.92 Å/pixel) from 2 datasets. 50,000 particles were selected from 2D classification (from dataset 1) and used for *ab initio* reconstruction to generate 5 classes. These were heterogeneously refined to generate two good classes (class 1 with 268,233 particles, and class 2 with 248,833 particles). These good particles were re-extracted with a box size of 440 pixels and a pixel size of 0.73 Å/pixel. The class 1 was refined with Non-uniform Refinement(*65*) and then Local Refinement to a resolution of 3.1 Å (U11 snRNP core map, M1). The signal subtraction approach was used to obtain a better-defined peripheral region of PDCD7. After particle subtraction, the reconstruction, which contains only PDCD7 region was low-pass filtered to 60 Å and used as a reference for 3D classification of 268,233 subtracted particles. 3D classification led to one good class, which contains 48,988 particles. This class was refined to 4.0 Å (map M4) and used to interpret the C-terminal leg (residues 388-485) of PDCD7^CC^ domain.

For the overall U11 snRNP, particles from class1 and class2 were merged together and refined with Non-uniform Refinement. This reconstruction was used as a reference to perform Heterogeneous refinement, which generated one good class. This class was refined with Non-uniform Refinement and then Local Refinement to a resolution of 3.4 Å (Overall U11 snRNP map, M2). Another Local Refinement was performed with a mask covering the Sm-region lobe of U11 snRNP, generating the U11 snRNP 3’-domain map 3 (M3) with a resolution of 3.0 Å. Particle subtraction was performed on these particles, and the reconstruction which contains only U11 snRNA SL4 region was low-pass filtered to 60 Å and used as a reference for 3D classification of 222,666 subtracted particles. 3D classification led to one good class, which contains 79,134 particles. This class was refined with Non-uniform Refinement and then Local Refinement to 3.7 Å (M5) and used to interpret the interaction between PDCD7(residues 171-255) and SNRNP48 (residues 7-32).

### Model building and validation

Templates for model building were generated by AlphaFold2(*41*). The templates were rigid-body fitted into the maps in ChimeraX(*66*). The models were then refined into the maps using ISOLDE(*67*). Subsequently, Coot was used manually adjust and build proteins and RNA(*68*). The stem-loop regions of U11 snRNA were generated in Coot and refined in ISOLDE, and the remaining nucleotides were added and real-space refined in Coot. Other components with poorly resolved densities (i.e. PDCD7^171-246^, SNRNP48) were docked into the map as rigid bodies and left in their original form. ZMAT5 and SNRNP48^Cterm^ binding sites were initially identified by an exhaustive *in silico* AlphaFold2-based search(*41*, *42*) for all possible interactions with Sm proteins using a previously described approach(*69*). A summary of the modelling is listed in the table S3. Final atomic models were refined to the high-resolution maps using Real-space Refinement in PHENIX(*70*) and validated using the wwPDB OneDep System(*71*). Additional validation statistics were obtained using MolProbity(*72*). Atomic models were visualized with ChimeraX or PyMol (Schrödinger).

---

**Figure S1.**
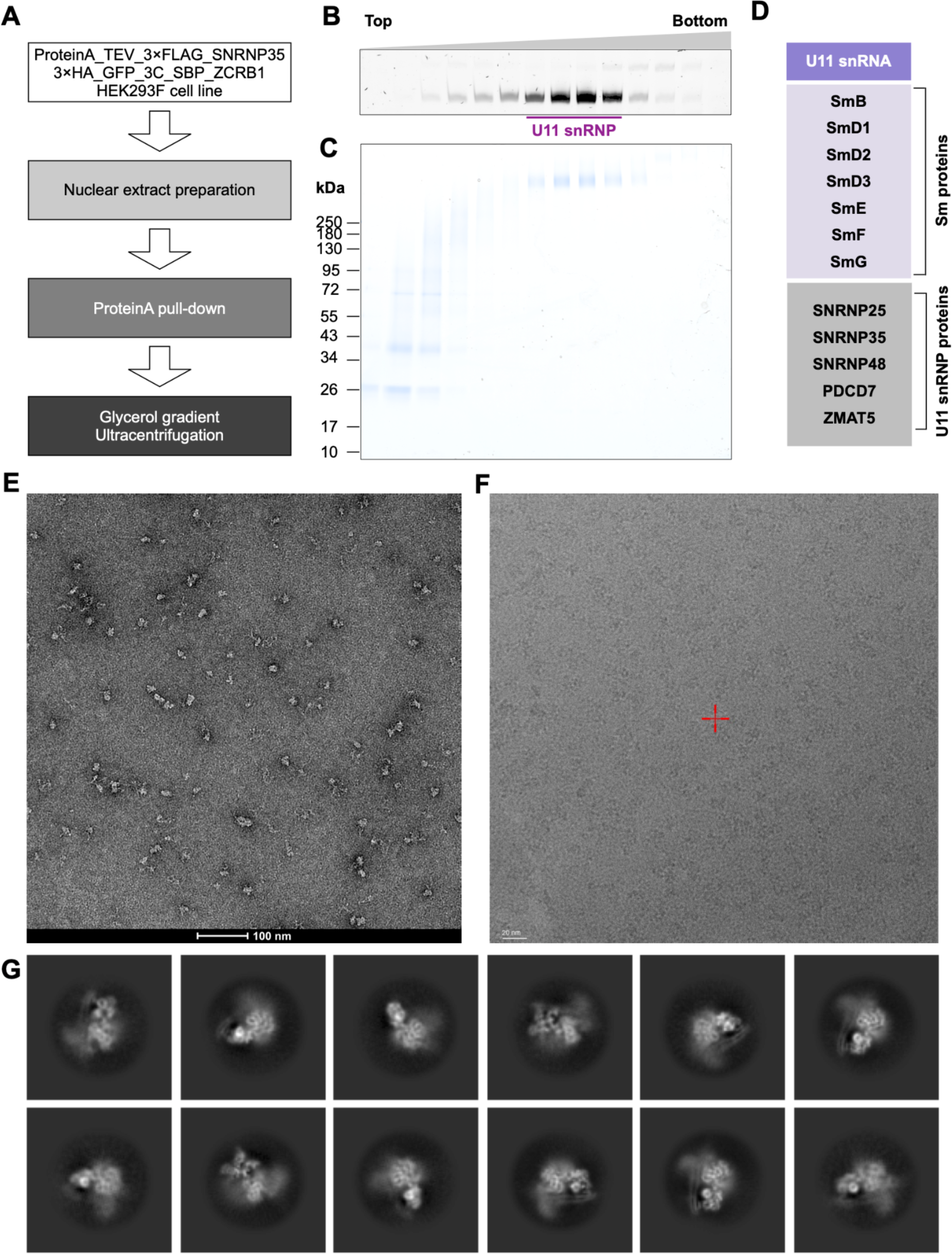
Isolation of U11 snRNP from HEK293F cells. (**A**) Experimental workflow used in sample purification. (**B**) UREA-PAGE analysis of the RNA extracted from native (no cross-linker) glycerol gradient ultracentrifugation. RNAs were stained with SYBR Gold and detected using Bio-Rad Chemidoc. (**C**) SDS-PAGE analysis of the fractions from the Grafix gradient. Proteins were stained with Coomassie blue. (**D**) List of proteins present in the U11 snRNP preparation. **(E)** A typical negative staining EM micrograph of the sample after Grafix gradient. (**F**) A representative cryo-EM micrograph of U11 snRNP sample collected on a 200kV Glacios/Falcon 4i microscope. (**G**) Selected 2D classes derived from cryo-EM data processing.

**Figure S2.**
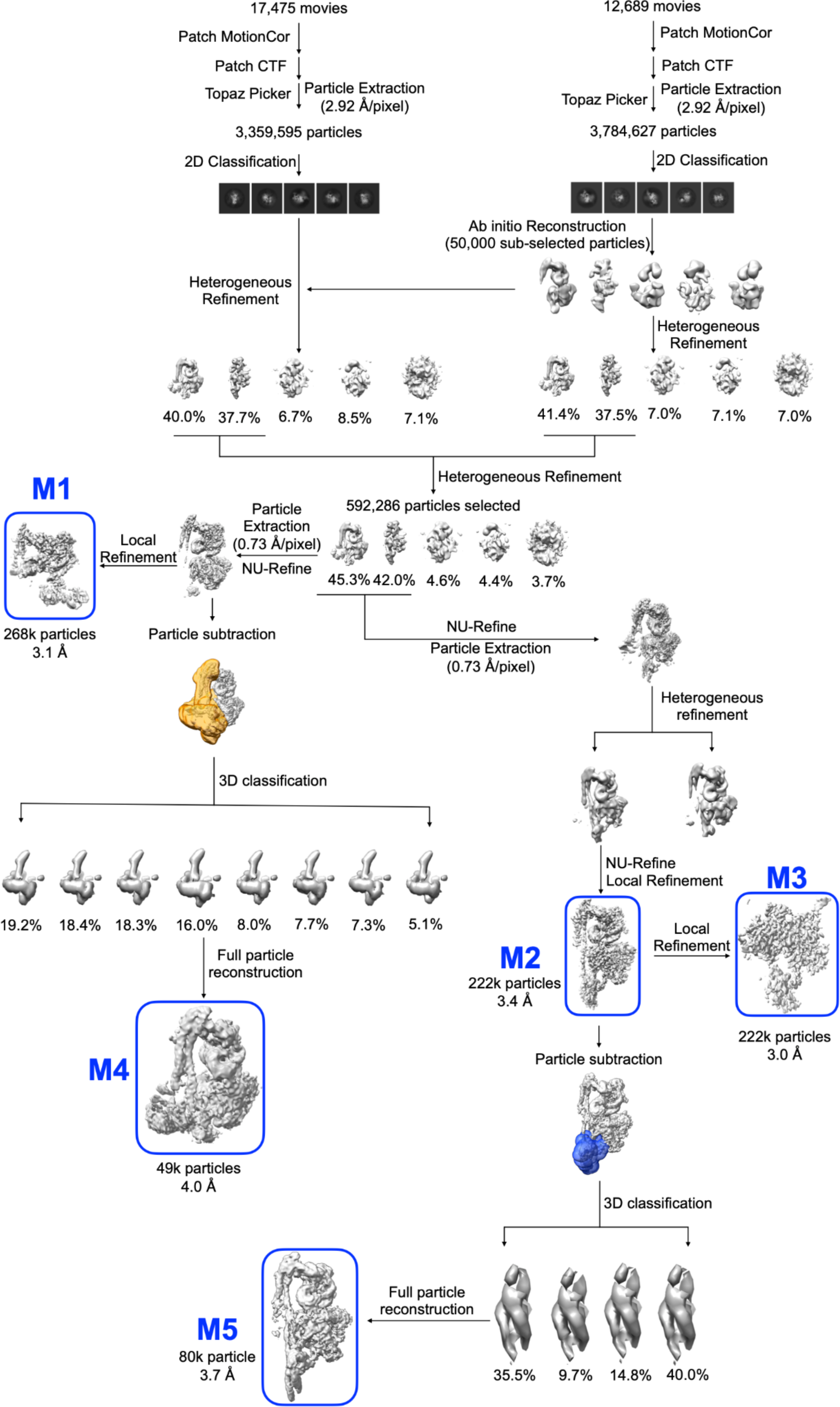
Cryo-EM data processing workflow.

**Figure S3.**
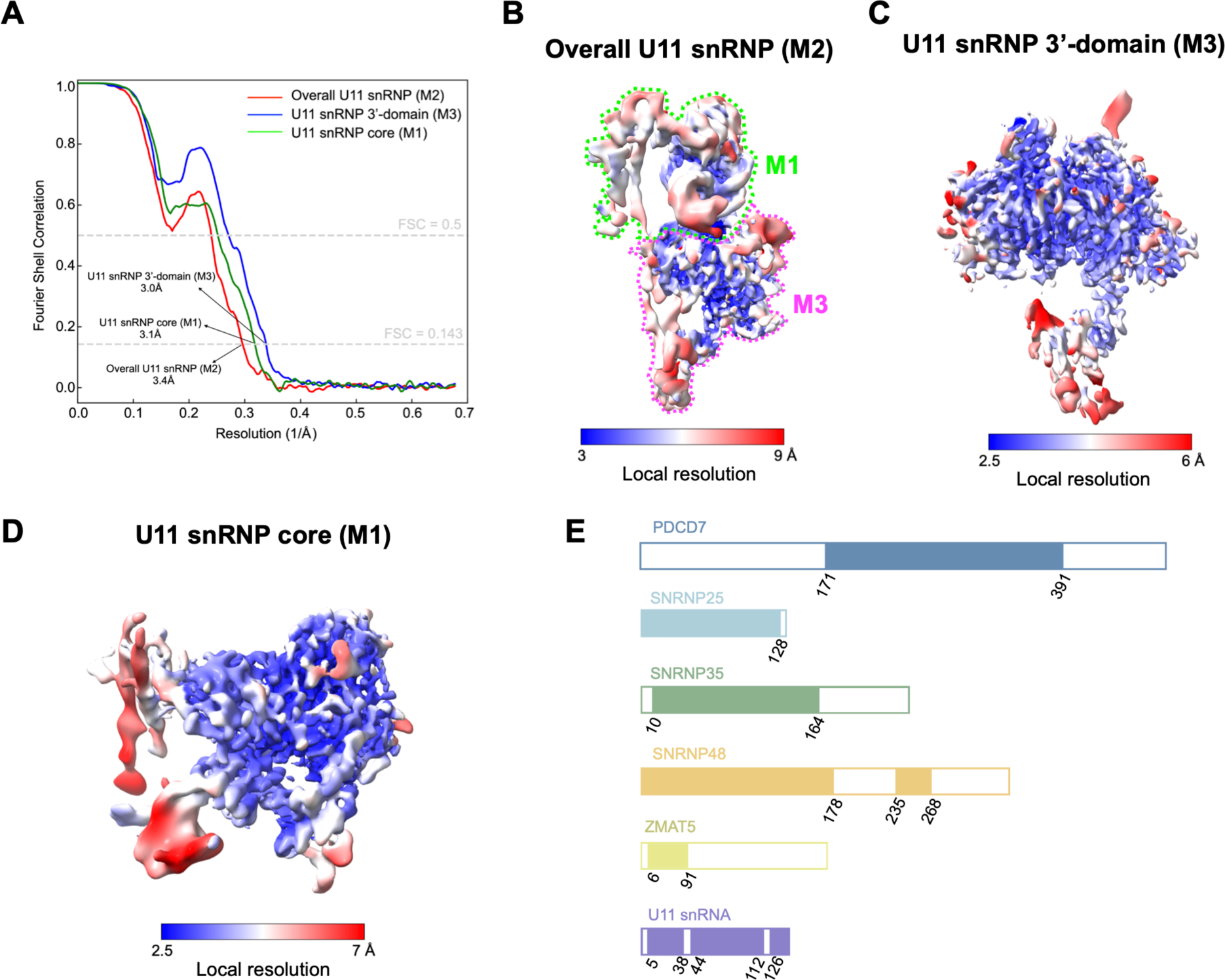
Global and local resolution analysis of the cryo-EM reconstructions. (**A**) Gold standard Fourier Shell Correlation (FSC) curves of masked maps determined in cryoSPARC v4.3.1. Red: Overall U11 snRNP; blue: U11 snRNP 3’-domain; green: U11 snRNP core. The local minima in the FSC curves are likely caused by the tight mask for local refinement. (**B**, **C**, **D**) Local resolution plotted on the isosurface of the locally filtered overall U11 snRNP map, U11 snRNP 3’-domain map and U11 snRNP core map, respectively. (**E**) Overview of the proteins modelled in the U11 snRNP structure. Filled-in rectangles denote the sequences that were modelled. Empty rectangles are parts of the proteins and RNA that could not be assigned to the reconstructed maps. Sm proteins are not shown here.

**Figure S4.**
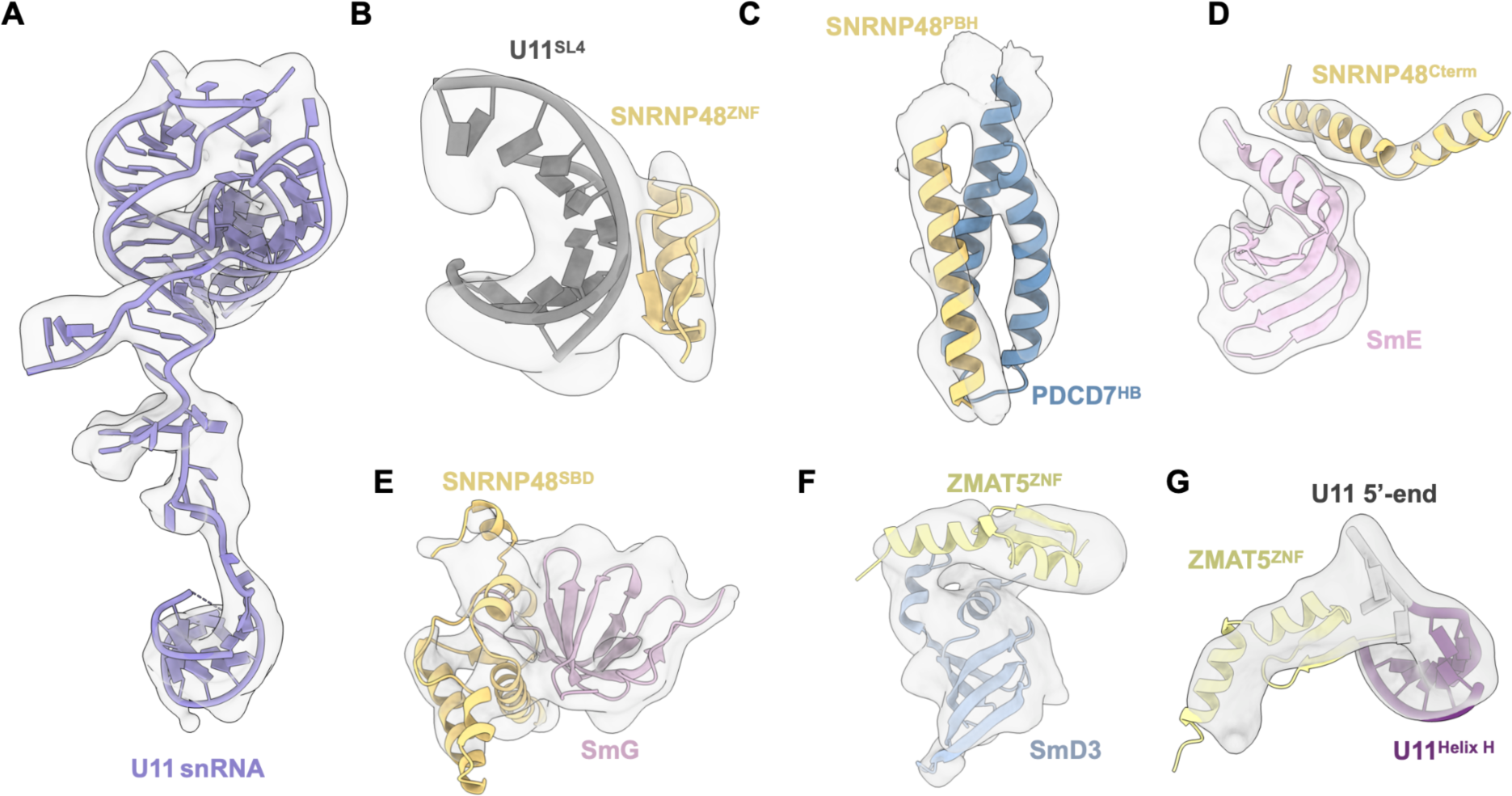
Low-resolution modelling of U11 snRNP. (**A**) Fitting of U11 snRNA into the U11 snRNP overall map M2. (**B**) Modelling of the SNRNP48 Znic finger domain, which binds to the minor groove of the U11^SL4^. (**C**) Fitting of three-helix bundle formed between PDCD7^HB^ and SNRNP48^PBH^. (**D**, **E**) Fitting of the SNRNP48 C-terminal helix (SNRNP48^Cterm^) and Sm binding domain (SNRNP48^SBD^) into the U11 snRNP 3’-domain map (M3), respectively. (**F**) The ZnC finger domain of ZMAT5 interacting with SmD3. (**G**) ZMAT5 is located directly next to 5’SS binding region of the U11 snRNA. PBH - PDCD7-Binding Helix; HB - Helix bundle; ZNF - Zinc Finger domain; SBD - Sm Binding Domain.Cterm - C-termianl helix.

**Figure S5.**
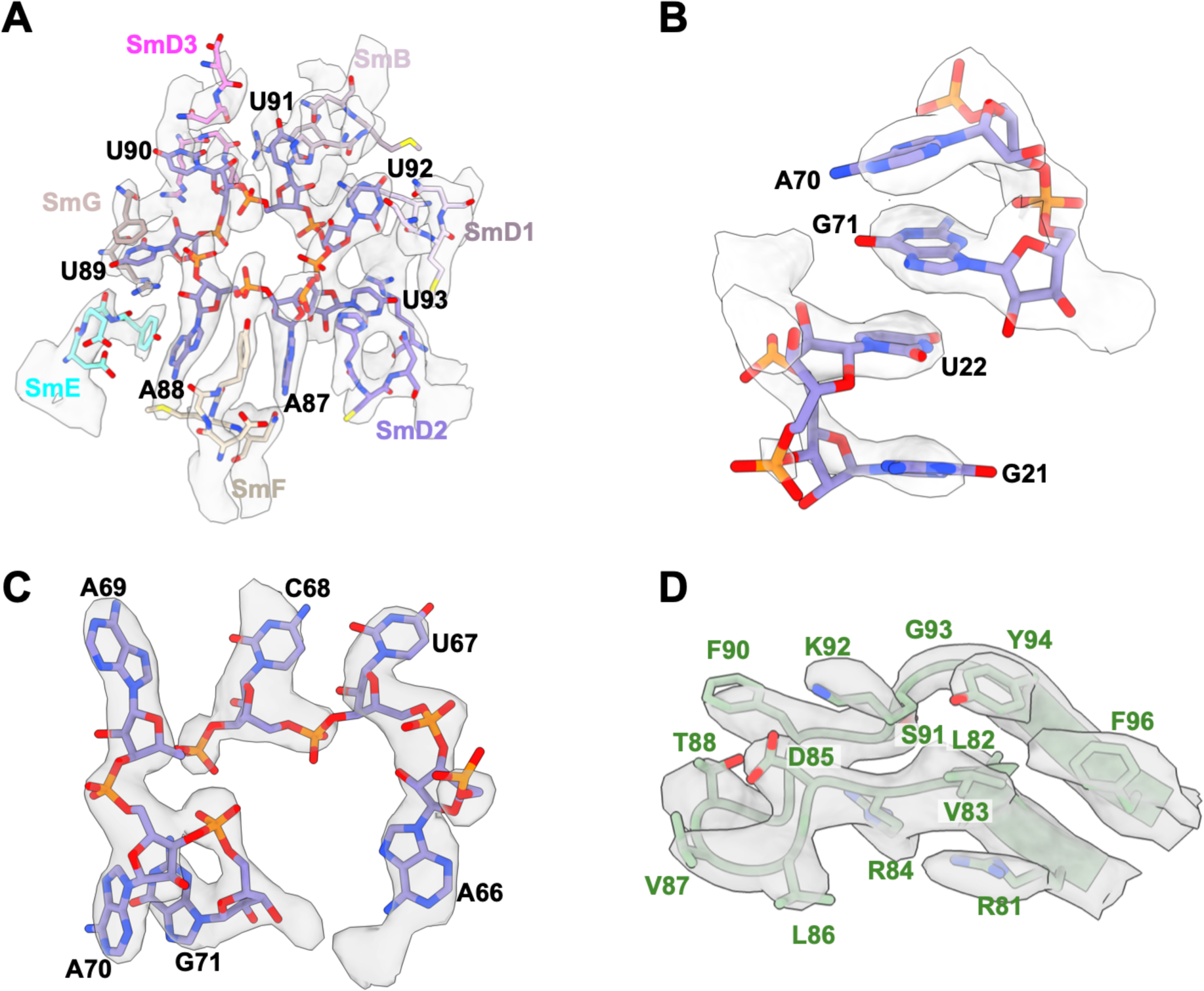
High-resolution modelling of U11 snRNP. (**A**) U11 snRNA Sm site (5’-AAUUUUU-3’) recognition at the central hole of the Sm ring (U11 snRNP 3’-domain map M3). (**B**) The atomic model of the bases located in the loops of SL1 and SL3 fitted into the U11 snRNP core map M1. (**C**) The atomic model of the SL3 loop fits into the U11 snRNP core map M1. (**D**) The atomic model of the loop extruding from the canonical SNRNP35^RRM^ domain.

**Figure S6:**
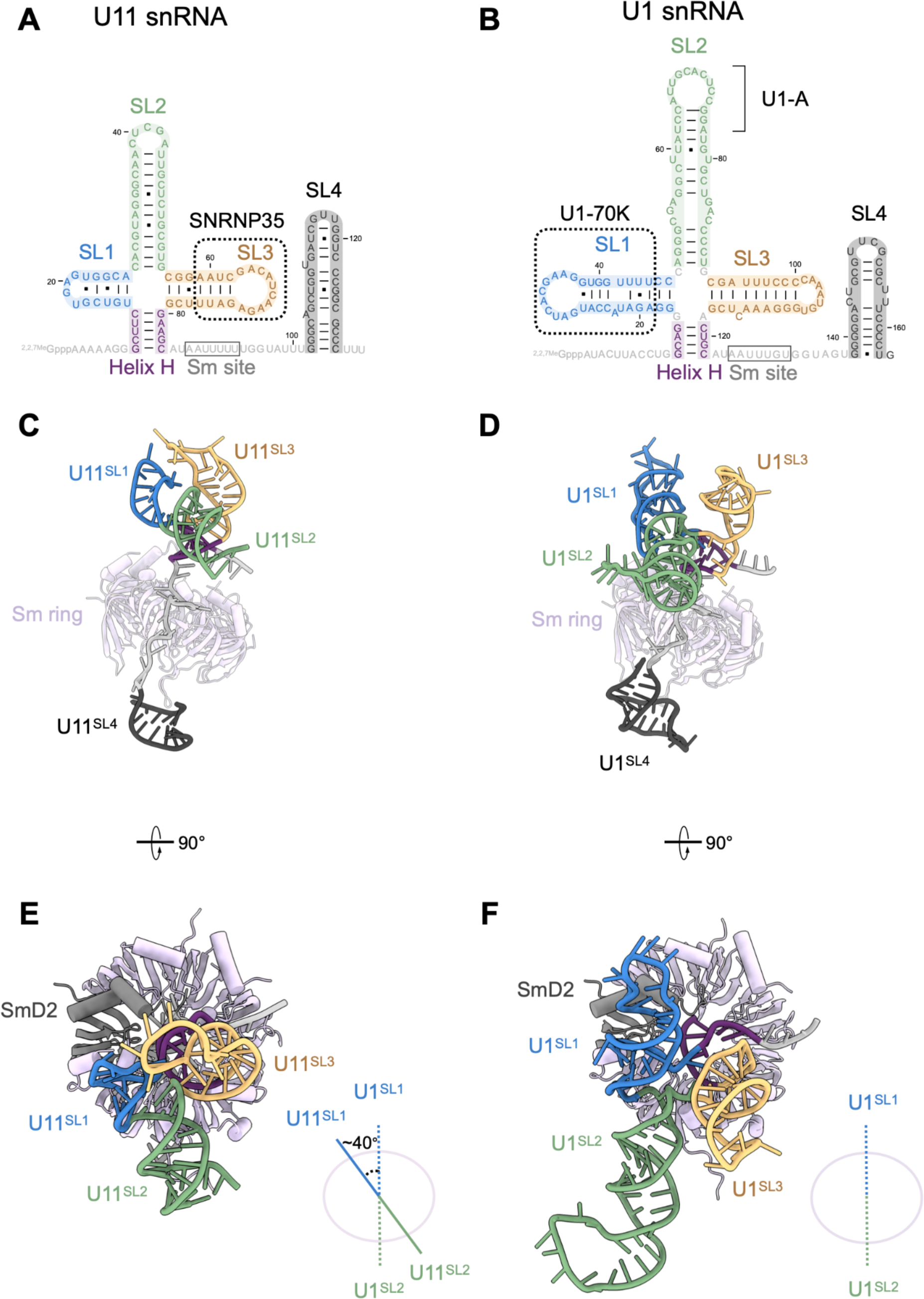
Structural comparison between U11 snRNA and U1 snRNA. (**A**) Secondary structure of U11 snRNA. The SL3 is coordinated by SNRNP35. (**B**) Secondary structure of the U1 snRNA. Based on the U1 snRNP structure (PDB: 4PJO), U1-70K and U1-A bind to SL1 and SL2 through their RRM domain, respectively. (**C**) Side view of the U11 snRNA showing the relative orientation of the stem-lops 1-4 with respect to the Sm ring (**D**) Side view of the U1 snRNA shown in the same orientation as U11 in (C), indicating the relative orientation of the stem-lops 1-4 with respect to the Sm ring. (**E**, **F**) Top view of the structures from (**C**) and (**D**), respectively, showing a relative rotation of the co-axially stacked stem loops SL1-SL2 with respect to the Sm ring. SmD2 (marked in grey) was used to align two models. Stem loops and Sm site are highlighted as shown in Figure 2.

**Figure S7.**
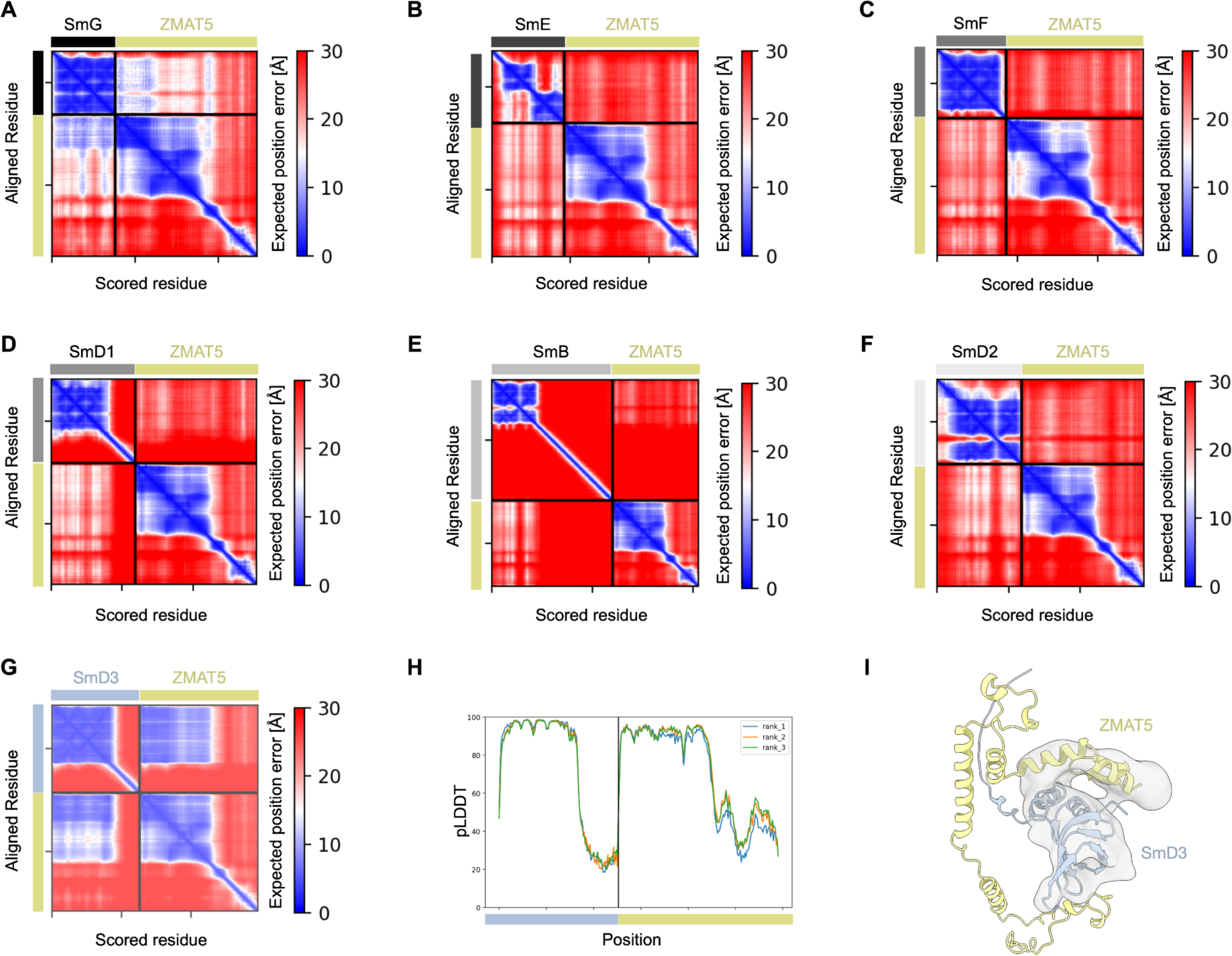
The AlphaFold2-based interaction screen between ZMAT5 and Sm proteins. (**A**-**G**) Predicted Aligned Error (PAE) plots of the ZMAT5:Sm proteins pairs. Cross-peaks with low PAE values indicated high confidence for the relative orientation of the regions corresponding to these cross-peaks. SmD3 was identified as the most likely interacting partner. (**H**) pLDDT score of the ZMAT5:SmD3 complex shown in (**G**) plotted against their sequences used for modelling. High pLDDT scores indicate high-confidence modelling as assessed by AlphaFold2 algorithm (*41*, *42*). (**I**) Fitting of the ZMAT5/SmD3 AlphaFold2 model into the U11 snRNP 3’-domain map (M3).

**Figure S8.**
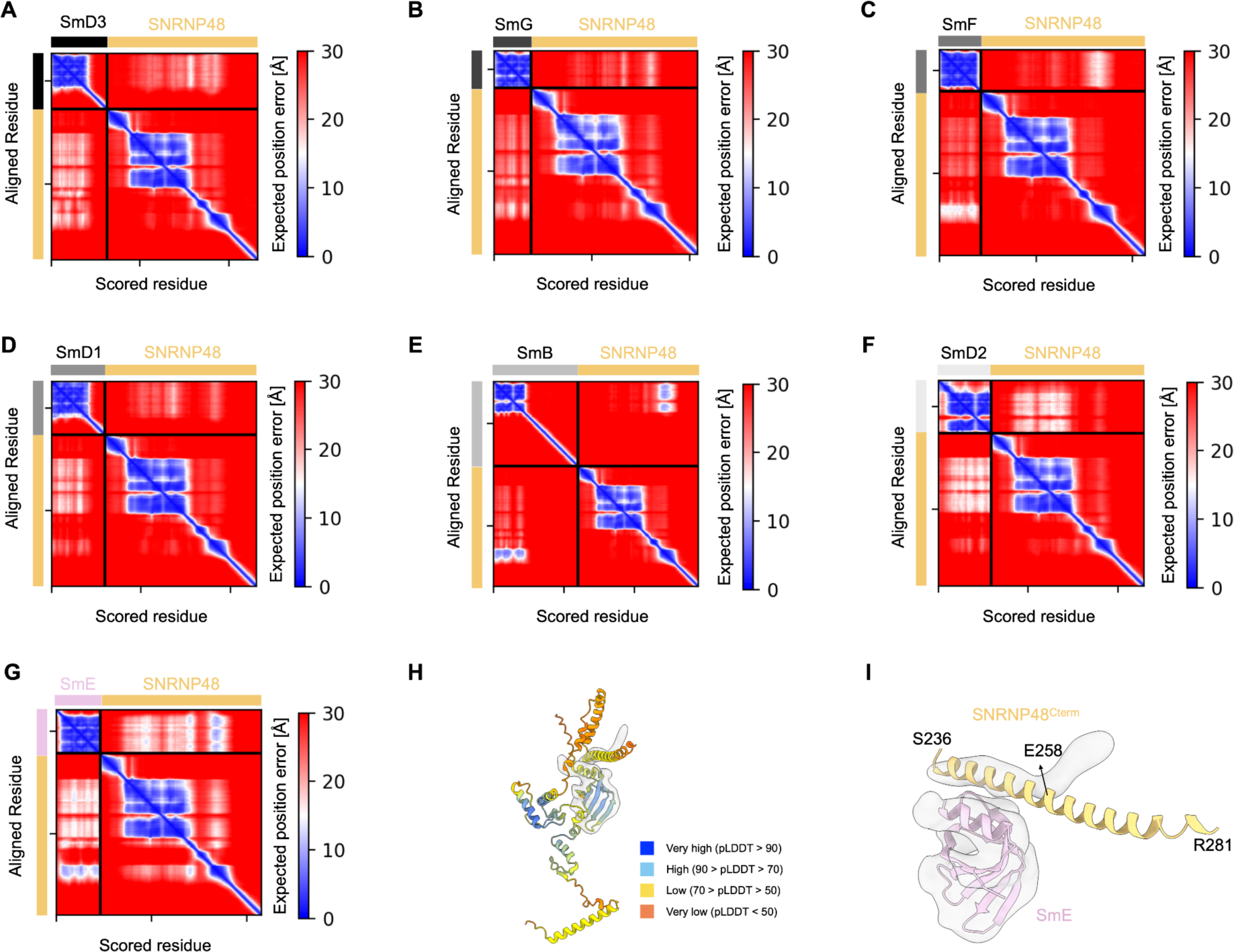
The AlphaFold2-based interaction screen between SNRNP48 and Sm proteins. (**A**-**G**) Predicted Aligned Error (PAE) plots of the complex formed by Sm proteins and SNRNP48. Cross-peaks with low PAE values indicated high confidence for the relative orientation of the regions corresponding to these cross-peaks. SmE was identified as the most likely interacting partner. (**H**) A cartoon representation of the SNRNP48/SmE AlphaFold2 model. Per-residue model confidence is assessed by AlphaFold2 algorithm (*41*, *42*) and coloured accordingly. (**I**) Close view of the SNRNP48/SmE AlphaFold2 model fitting to U11 snRNP 3’-domain map (M3). The first half of the helix (236-258) fits well into the map, and the second half (258-281) needs to be adjusted accordingly. Cterm - C-termianl helix.

**Figure S9.**
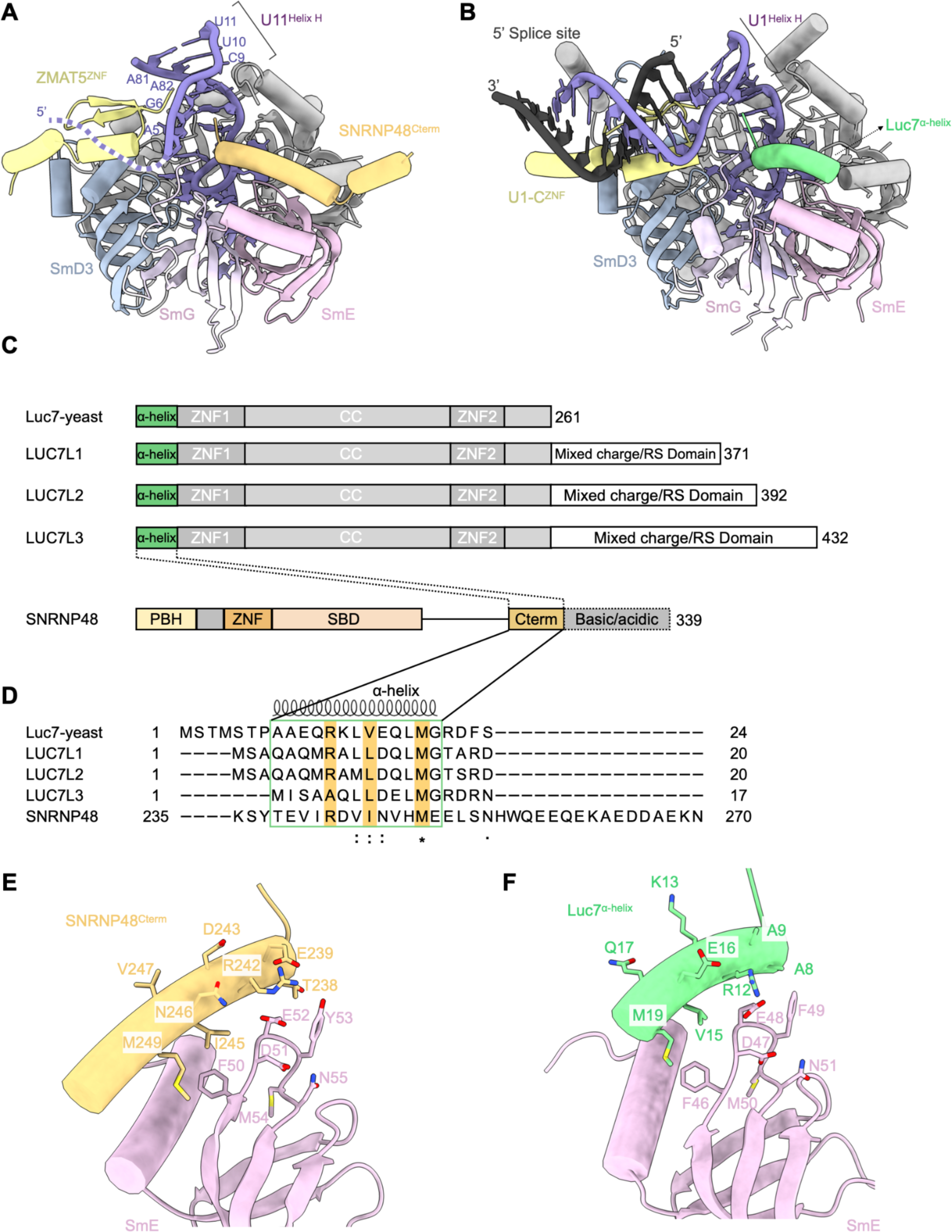
5’SS binding region of human U11 snRNA and yeast U1 snRNA. (**A**) 5’SS binding region of human U11 snRNA is positioned between ZMAT5^ZNF^ and SNRNP48^Cterm^ domains, suggesting their possible involvement in the stabilisation of the 5’SS binding. (**B**) An equivalent view of the yeast U1 snRNP bound to the model 5’SS (*37*). The α-helix of Luc7 interacts with 5’SS/U1 duplex, which is similar to the role of SNRNP48^Cterm^ domain in human U11 snRNP. (**C**) Domain architectures of yeast Luc7 protein, three human LUC7 like proteins and SNRNP48; PBH - PDCD7-Binding Helix; CC - coiled coil; ZNF - Zinc Finger domain; SBD - Sm Binding Domain.Cterm - C-termianl helix. (**D**) Sequence alignment of the proteins shown in (**C**). The C-terminal helix of SNRNP48 is used to align with the first α-helix of Luc7 and LUC7-like proteins. Three conserved residues, as indicated in (**E**, **F**), are coloured in brown. (**E**) Close view of the interactions between SNRNP48^Cterm^ and SmE based on AF2 modelling and fitting to the low-resolution cryo-EM map. (**F**) Close view of the interactions between the α-helix of Luc7 and SmE. Structurally conserved residues are indicated in (**D**).

**Figure S10.**
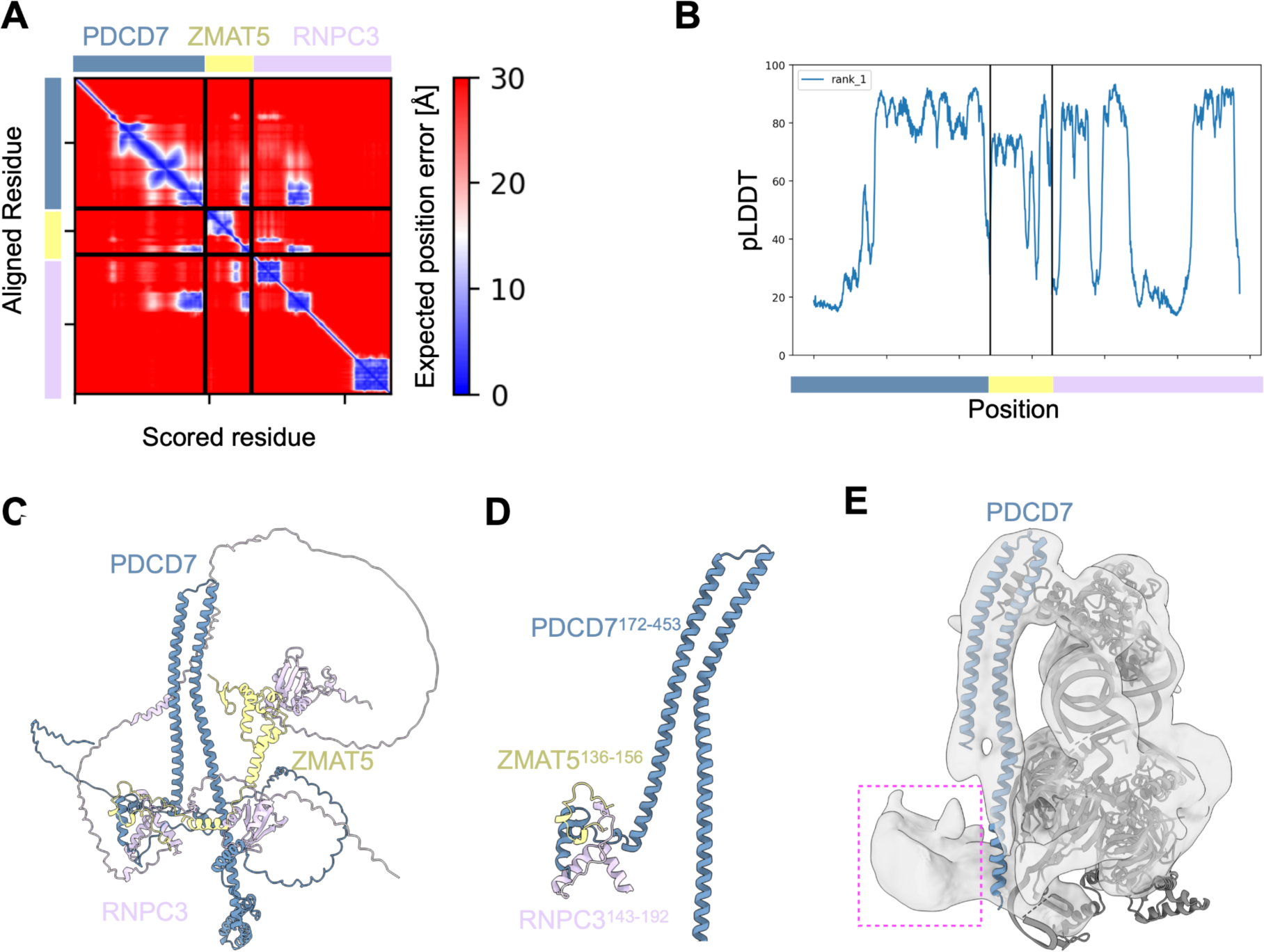
AlphaFold2-based modelling of the interaction between PDCD7, ZMAT5 and RNPC3. (**A**) Predicted Aligned Error (PAE) plot of the complex PDCD7/ZMAT5/RNPC3. Cross-peaks with low PAE values indicated high confidence for the relative orientation of the regions corresponding to these cross-peaks. (**B**) pLDDT score of the complex plotted against the sequences used for modelling. High pLDDT scores indicate high-confidence modelling as assessed by the AlphaFold2 algorithm(*41*, *42*). (**C**) Cartoon representation of the PDCD7/ZMAT5/RNPC3 AF2 model. (**D**) A clear view of the interaction among PDCD7^172-453^, ZMAT5^136-156^ and RNPC3^143-192^. (**E**) A low-pass filtered map of the U11 snRNP (M4) with the U11 snRNP structure fitted in. The fuzzy density marked by a magenta square, overlaps with the AlphaFold2 model of these three proteins superimposed on the experimental structure.

**Table S1.** Solution mass spectrometry analysis of the purified U11 snRNP.

| No<sup>1</sup> | Protein | Unique peptides | Total spectral count | Total Intensity |
| --- | --- | --- | --- | --- |
| 2 | RNPC3 | 43 | 67 | 3,41E+09 |
| 3 | PDCD7 | 28 | 38 | 1,27E+09 |
| 4 | SNRNP48 | 23 | 29 | 1,06E+09 |
| 5 | SNRNP25 | 6 | 6 | 6,46E+08 |
| 6 | SNRNP35 | 22 | 24 | 4,81E+08 |
| 15 | SNRPD2 | 13 | 20 | 2,64E+08 |
| 24 | SNRPD3 | 4 | 4 | 1,44E+08 |
| 28 | ZMAT5 | 9 | 11 | 9,79E+07 |
| 32 | SNRPG | 1 | 7 | 7,73E+07 |
| 33 | SNRPB | 0 | 6 | 6,61E+07 |
| 34 | SF3B3 | 20 | 24 | 6,41E+07 |
| 37 | SF3B1 | 31 | 36 | 5,63E+07 |
| 52 | SNRPD1 | 5 | 5 | 3,05E+07 |
| 59 | SNRPE | 2 | 3 | 2,52E+07 |
| 74 | SF3B2 | 19 | 27 | 1,15E+07 |

<sup>1</sup>Position on the hits list, when sorted by Total intensity in descending order.

**Table S2.** Cryo-EM data collection, refinement and validation statistics.

|  | U11 snRNP core | Overall U11 snRNP | U11 snRNP 3'-domain |
| --- | --- | --- | --- |
| **Data collection and processing** |  |  |  |
| Microscope | TFS Krios |  |  |
| Voltage (keV) | 300 |  |  |
| Camera | Falcon 4i |  |  |
| Magnification | 165kx |  |  |
| Pixel size at detector (Å/pixel) | 0.73 |  |  |
| Total electron exposure (e<sup>-</sup>/Å<sup>2</sup>)<br>(dataset1/dataset2) | 41.73/42.61 |  |  |
| Defocus range (µm)<br>(dataset1/dataset2) | -0.8 to -1.8/-0.7 to -1.7 |  |  |
| Automation software | SerialEM |  |  |
| Energy filter slit width | 20 eV |  |  |
| Micrographs collected (no.) | 31801 |  |  |
| Micrographs used (no.) | 30164 |  |  |
| Total extracted particles (no.) | 7,144,222 |  |  |
| **For each reconstruction:** | **M1** | **M2** | **M3** |
| Final particles (no.) | 268,233 | 222,666 | 222,666 |
| Point-group | C1 | C1 | C1 |
| Resolution (global, Å) |  |  |  |
| FSC 0.5 (unmasked/masked) | 3.40/3.50 | 4.02/4.14 | 3.44/3.36 |
| FSC 0.143 (unmasked/masked) | 3.09/3.14 | 3.38/3.46 | 2.96/2.93 |
| Map sharpening B factor (Å<sup>2</sup>)<br>(main maps are unsharpened) | -75.9 | -71.2 | -55.4 |
| **Model composition** |  |  |  |
| Protein (aa) | 377 | 1324 | 784 |
| RNA (bases) | 66 | 109 | 43 |
| **Model refinement** |  |  |  |
| Refinement package | Phenix | Phenix | Phenix |
| real or reciprocal space | real | real | real |
| resolution cutoff (Å) | 3.1 | 3.4 | 3.0 |
| CC (mask) | 0.78 | 0.68 | 0.70 |
| B factors (Å<sup>2</sup>) |  |  |  |
| Protein residues | 142.82 | 167.72 | 100.41 |
| RNA | 123.19 | 178.38 | 153.40 |
| R.m.s d from ideal values |  |  |  |
| Bond lengths (Å) | 0.005 | 0.004 | 0.004 |
| Bond angles (°) | 0.818 | 0.859 | 0.914 |
| **Validation** |  |  |  |
| MolProbity score | 2.02 | 2.23 | 2.41 |
| CaBLAM outliers | 1.37 | 3.23 | 3.38 |
| Clashscore | 18.54 | 18.60 | 15.39 |
| Rotamers outliers (%) | 0.93 | 3.23 | 2.10 |
| C-beta outliers (%) | 0.00 | 0.00 | 0.00 |
| EMRinger score | 2.49 | 1.22 | 1.92 |
| Ramachandran plot |  |  |  |
| Favored (%) | 96.23 | 92.67 | 92.26 |
| Outliers (%) | 0.54 | 0.15 | 0.00 |

**Table S3.** Summary of the protein and RNA modelling.

| Overall U11 snRNP |  |  |  |  |  |  |
| --- | --- | --- | --- | --- | --- | --- |
| Molecule | Chain ID | Uniprot ID | Total Res. | Modelled Res. | Template used | Modelling approach |
| U11 snRNA | A |  | 135 | 5-38, 44-112, 126-131 |  | De novo & docked |
| SNRNP25 | B | Q9BV90 | 132 | 1-128 | AlphaFold | Docked & adjusted |
| SNRNP35 | C | Q16560 | 246 | 10-164 | AlphaFold | Docked & adjusted |
| PDCD7 | D | Q8N8D1 | 485 | 171-391 | AlphaFold | Docked & adjusted |
| ZMAT5 | E | Q9UDW3 | 170 | 2-43 | AlphaFold | Docked |
| SNRNP48 | F | Q6IEG0 | 339 | 1-178, 235-268 | AlphaFold | Docked & adjusted |
| SmB | k | P14678 | 240 | 6-91 | 4PJO | Docked & adjusted |
| SmD1 | h | P62314 | 119 | 2-82 | 4PJO | Docked & adjusted |
| SmD2 | i | P62316 | 118 | 11-114 | 4PJO | Docked & adjusted |
| SmD3 | j | P62318 | 126 | 3-83 | 4PJO | Docked & adjusted |
| SmE | l | P62304 | 92 | 14-90 | 4PJO | Docked & adjusted |
| SmF | m | P62306 | 86 | 2-75 | 4PJO | Docked & adjusted |
| SmG | n | P62308 | 76 | 4-76 | 4PJO | Docked & adjusted |

## Notes

### Competing Interest Statement

The authors have declared no competing interest.

## References and Notes

1. C. L. Will, R. Lührmann, Spliceosome structure and function. Cold Spring Harb. Perspect. Biol. 3 (2011), doi:10.1101/cshperspect.a003707.

2. A. G. Matera, Z. Wang, A day in the life of the spliceosome. Nat. Rev. Mol. Cell Biol. 15, 108– 121 (2014).

3. P. Papasaikas, J. Valcárcel, The Spliceosome: The Ultimate RNA Chaperone and Sculptor. Trends Biochem. Sci. 41, 33–45 (2016).

4. W.-Y. Tarn, J. A. Steitz, A Novel Spliceosome Containing U11, U12, and U5 snRNPs Excises a Minor Class (AT–AC) Intron In Vitro. Cell. 84, 801–811 (1996).

5. S. L. Hall, R. A. Padgett, Requirement of U12 snRNA for in vivo splicing of a minor class of eukaryotic nuclear pre-mRNA introns. Science. 271, 1716–1718 (1996).

6. C. L. Will, C. Schneider, M. Hossbach, H. Urlaub, R. Rauhut, S. Elbashir, T. Tuschl, R. Lührmann, The human 18S U11/U12 snRNP contains a set of novel proteins not found in the U2-dependent spliceosome. RNA. 10, 929–941 (2004).

7. R. Bai, R. Wan, L. Wang, K. Xu, Q. Zhang, J. Lei, Y. Shi, Structure of the activated human minor spliceosome. Science. 371 (2021), doi:10.1126/science.abg0879.

8. B. de Wolf, A. Oghabian, M. V. Akinyi, S. Hanks, E. C. Tromer, J. J. E. van Hooff, L. van Voorthuijsen, L. E. van Rooijen, J. Verbeeren, E. C. H. Uijttewaal, M. P. A. Baltissen, S. Yost, P. Piloquet, M. Vermeulen, B. Snel, B. Isidor, N. Rahman, M. J. Frilander, G. J. P. L. Kops, Chromosomal instability by mutations in the novel minor spliceosome component CENATAC. EMBO J. 40, e106536 (2021).

9. T. Suzuki, T. Shinagawa, T. Niwa, H. Akeda, S. Hashimoto, H. Tanaka, Y. Hiroaki, F. Yamasaki, H. Mishima, T. Kawai, T. Higashiyama, K. Nakamura, The DROL1 subunit of U5 snRNP in the spliceosome is specifically required to splice AT-AC-type introns in Arabidopsis. Plant J. 109, 633–648 (2022).

10. A. J. Norppa, I. Chowdhury, L. E. van Rooijen, J. J. Ravantti, B. Snel, M. Varjosalo, M. J. Frilander, Distinct functions for the paralogous RBM41 and U11/U12-65K proteins in the

12. A. G. Russell, J. M. Charette, D. F. Spencer, M. W. Gray, An early evolutionary origin for the minor spliceosome. Nature. 443, 863–866 (2006).

13. G. E. Larue, S. W. Roy, Where the minor things are: a pan-eukaryotic survey suggests neutral processes may explain much of minor intron evolution. Nucleic Acids Res. (2023), doi:10.1093/nar/gkad797.

14. B. Verma, M. V. Akinyi, A. J. Norppa, M. J. Frilander, Minor spliceosome and disease. Semin. Cell Dev. Biol. 79, 103–112 (2018).

15. J. J. Turunen, E. H. Niemelä, B. Verma, M. J. Frilander, The significant other: splicing by the minor spliceosome. Wiley Interdiscip. Rev. RNA. 4, 61–76 (2013).

16. C. B. Burge, R. A. Padgett, P. A. Sharp, Evolutionary fates and origins of U12-type introns. Mol. Cell. 2, 773–785 (1998).

17. A. A. Hoskins, L. J. Friedman, S. S. Gallagher, D. J. Crawford, E. G. Anderson, R. Wombacher, N. Ramirez, V. W. Cornish, J. Gelles, M. J. Moore, Ordered and dynamic assembly of single spliceosomes. Science. 331, 1289–1295 (2011).

18. R. Das, Z. Zhou, R. Reed, Functional association of U2 snRNP with the ATP-independent spliceosomal complex E. Mol. Cell. 5, 779–787 (2000).

19. M. M. Konarska, P. A. Sharp, Electrophoretic separation of complexes involved in the splicing of precursors to mRNAs. Cell. 46, 845–855 (1986).

20. C. M. Newnham, C. C. Query, The ATP requirement for U2 snRNP addition is linked to the pre-mRNA region 5’ to the branch site. RNA. 7, 1298–1309 (2001).

21. K. M. Wassarman, J. A. Steitz, The low-abundance U11 and U12 small nuclear ribonucleoproteins (snRNPs) interact to form a two-snRNP complex. Mol. Cell. Biol. 12, 1276–1285 (1992).

22. M. J. Frilander, J. A. Steitz, Initial recognition of U12-dependent introns requires both U11/5’ splice-site and U12/branchpoint interactions. Genes Dev. 13, 851–863 (1999).

23. C. L. Will, C. Schneider, R. Reed, R. Lührmann, Identification of both shared and distinct proteins in the major and minor spliceosomes. Science. 284, 2003–2005 (1999).

24. H. Shen, X. Zheng, S. Luecke, M. R. Green, The U2AF35-related protein Urp contacts the 3’ splice site to promote U12-type intron splicing and the second step of U2-type intron splicing. Genes Dev. 24, 2389–2394 (2010).

25. H. Benecke, R. Lührmann, C. L. Will, The U11/U12 snRNP 65K protein acts as a molecular bridge, binding the U12 snRNA and U11-59K protein. EMBO J. 24, 3057–3069 (2005).

26. J. J. Turunen, C. L. Will, M. Grote, R. Lührmann, M. J. Frilander, The U11-48K protein contacts the 5’ splice site of U12-type introns and the U11-59K protein. Mol. Cell. Biol. 28, 3548–3560 (2008).

27. H. Tidow, A. Andreeva, T. J. Rutherford, A. R. Fersht, Solution structure of the U11-48K CHHC zinc-finger domain that specifically binds the 5’ splice site of U12-type introns. Structure. 17, 294–302 (2009).

28. B. Kastner, C. L. Will, H. Stark, R. Lührmann, Structural Insights into Nuclear pre-mRNA Splicing in Higher Eukaryotes. Cold Spring Harb. Perspect. Biol. 11 (2019), doi:10.1101/cshperspect.a032417.

29. M. E. Wilkinson, C. Charenton, K. Nagai, RNA Splicing by the Spliceosome. Annu. Rev. Biochem. 89, 359–388 (2020).

30. L. Zhang, A. Vielle, S. Espinosa, R. Zhao, RNAs in the spliceosome: Insight from cryoEM structures. Wiley Interdiscip. Rev. RNA. 10, e1523 (2019).

31. R. Wan, R. Bai, X. Zhan, Y. Shi, How Is Precursor Messenger RNA Spliced by the Spliceosome? Annu. Rev. Biochem. 89, 333–358 (2020).

32. J. Tholen, W. P. Galej, Structural studies of the spliceosome: Bridging the gaps. Curr. Opin. Struct. Biol. 77, 102461 (2022).

33. D. A. Pomeranz Krummel, C. Oubridge, A. K. W. Leung, J. Li, K. Nagai, Crystal structure of human spliceosomal U1 snRNP at 5.5 A resolution. Nature. 458, 475–480 (2009).

34. G. Weber, S. Trowitzsch, B. Kastner, R. Lührmann, M. C. Wahl, Functional organization of the Sm core in the crystal structure of human U1 snRNP. EMBO J. 29, 4172–4184 (2010).

35. X. Li, S. Liu, J. Jiang, L. Zhang, S. Espinosa, R. C. Hill, K. C. Hansen, Z. H. Zhou, R. Zhao, CryoEM structure of Saccharomyces cerevisiae U1 snRNP offers insight into alternative splicing. Nat. Commun. 8, 1035 (2017).

36. Y. Kondo, C. Oubridge, A.-M. M. van Roon, K. Nagai, Crystal structure of human U1 snRNP, a small nuclear ribonucleoprotein particle, reveals the mechanism of 5’ splice site recognition. Elife. 4 (2015), doi:10.7554/eLife.04986.

37. C. Plaschka, P.-C. Lin, C. Charenton, K. Nagai, Prespliceosome structure provides insights into spliceosome assembly and regulation. Nature. 559, 419–422 (2018).

38. C. C. Query, R. C. Bentley, J. D. Keene, A common RNA recognition motif identified within a defined U1 RNA binding domain of the 70K U1 snRNP protein. Cell. 57, 89–101 (1989).

39. C. Oubridge, N. Ito, P. R. Evans, C. H. Teo, K. Nagai, Crystal structure at 1.92 A resolution of the RNA-binding domain of the U1A spliceosomal protein complexed with an RNA hairpin. Nature. 372, 432–438 (1994).

40. G. M. Daubner, A. Cléry, F. H.-T. Allain, RRM-RNA recognition: NMR or crystallography…and new findings. Curr. Opin. Struct. Biol. 23, 100–108 (2013).

41. J. Jumper, R. Evans, A. Pritzel, T. Green, M. Figurnov, O. Ronneberger, K. Tunyasuvunakool, R. Bates, A. Žídek, A. Potapenko, A. Bridgland, C. Meyer, S. A. A. Kohl, A. J. Ballard, A. Cowie, B. Romera-Paredes, S. Nikolov, R. Jain, J. Adler, T. Back, S. Petersen, D. Reiman, E. Clancy, M. Zielinski, M. Steinegger, M. Pacholska, T. Berghammer, S. Bodenstein, D. Silver, O. Vinyals, A. W. Senior, K. Kavukcuoglu, P. Kohli, D. Hassabis, Highly accurate protein structure prediction with AlphaFold. Nature. 596, 583–589 (2021).

42. M. Mirdita, K. Schütze, Y. Moriwaki, L. Heo, S. Ovchinnikov, M. Steinegger, ColabFold: making protein folding accessible to all. Nat. Methods. 19, 679–682 (2022).

43. A. J. Norppa, T. M. Kauppala, H. A. Heikkinen, B. Verma, H. Iwaï, M. J. Frilander, Mutations in the U11/U12-65K protein associated with isolated growth hormone deficiency lead to structural destabilization and impaired binding of U12 snRNA. RNA. 24, 396–409 (2018).

44. R. L. Nelissen, C. L. Will, W. J. van Venrooij, R. Lührmann, The association of the U1-specific 70K and C proteins with U1 snRNPs is mediated in part by common U snRNP proteins. EMBO J. 13, 4113–4125 (1994).

45. I. Kolossova, R. A. Padgett, U11 snRNA interacts in vivo with the 5’ splice site of U12-dependent (AU-AC) pre-mRNA introns. RNA. 3, 227–233 (1997).

46. R. C. Dietrich, J. D. Fuller, R. A. Padgett, A mutational analysis of U12-dependent splice site dinucleotides. RNA. 11, 1430–1440 (2005).

47. X. Roca, A. R. Krainer, I. C. Eperon, Pick one, but be quick: 5’ splice sites and the problems of too many choices. Genes Dev. 27, 129–144 (2013).

48. Y.-T. Yu, J. A. Steitz, Site-specific crosslinking of mammalian U11 and U6atac to the 5ʹ splice site of an AT–AC intron. Proceedings of the National Academy of Sciences. 94, 6030–6035 (1997).

49. M. E. Wilkinson, S. M. Fica, W. P. Galej, C. M. Norman, Postcatalytic spliceosome structure reveals mechanism of 3ʹ–splice site selection. Science (2017) (available at https://www.science.org/doi/abs/10.1126/science.aar3729).

50. O. Puig, E. Bragado-Nilsson, T. Koski, B. Séraphin, The U1 snRNP-associated factor Luc7p affects 5’ splice site selection in yeast and human. Nucleic Acids Res. 35, 5874–5885 (2007).

51. X. Li, S. Liu, L. Zhang, A. Issaian, R. C. Hill, S. Espinosa, S. Shi, Y. Cui, K. Kappel, R. Das, K. C. Hansen, Z. H. Zhou, R. Zhao, A unified mechanism for intron and exon definition and back-splicing. Nature. 573, 375–380 (2019).

52. N. J. Daniels, C. E. Hershberger, X. Gu, C. Schueger, W. M. DiPasquale, J. Brick, Y. Saunthararajah, J. P. Maciejewski, R. A. Padgett, Functional analyses of human LUC7-like proteins involved in splicing regulation and myeloid neoplasms. Cell Rep. 35, 108989 (2021).

53. A. A. Jourdain, B. E. Begg, E. Mick, H. Shah, S. E. Calvo, O. S. Skinner, R. Sharma, S. M. Blue, G. W. Yeo, C. B. Burge, V. K. Mootha, Loss of LUC7L2 and U1 snRNP subunits shifts energy metabolism from glycolysis to OXPHOS. Mol. Cell. 81, 1905–1919.e12 (2021).

54. C. J. Kenny, M. P. McGurk, C. B. Burge, Human LUC7 proteins impact splicing of two major subclasses of 5’ splice sites. bioRxiv (2022), doi:10.1101/2022.12.07.519539.

55. S. Zhang, S. Aibara, S. M. Vos, D. E. Agafonov, R. Lührmann, P. Cramer, Structure of a transcribing RNA polymerase II–U1 snRNP complex. Science. 371, 305–309 (2021).

56. J. D. Dignam, R. M. Lebovitz, R. G. Roeder, Accurate transcription initiation by RNA polymerase II in a soluble extract from isolated mammalian nuclei. Nucleic Acids Res. 11, 1475–1489 (1983).

57. T. W. Nilsen, Preparation of Nuclear Extracts from HeLa cells. Cold Spring Harb. Protoc. 2013, 579–583 (2013).

58. B. Kastner, N. Fischer, M. M. Golas, B. Sander, P. Dube, D. Boehringer, K. Hartmuth, J. Deckert, F. Hauer, E. Wolf, H. Uchtenhagen, H. Urlaub, F. Herzog, J. M. Peters, D. Poerschke, R. Lührmann, H. Stark, GraFix: sample preparation for single-particle electron cryomicroscopy. Nat. Methods. 5, 53–55 (2008).

59. C. S. Hughes, S. Foehr, D. A. Garfield, E. E. Furlong, L. M. Steinmetz, J. Krijgsveld, Ultrasensitive proteome analysis using paramagnetic bead technology. Mol. Syst. Biol. 10, 757 (2014).

60. H. Franken, T. Mathieson, D. Childs, G. M. A. Sweetman, T. Werner, I. Tögel, C. Doce, S. Gade, M. Bantscheff, G. Drewes, F. B. M. Reinhard, W. Huber, M. M. Savitski, Thermal proteome profiling for unbiased identification of direct and indirect drug targets using multiplexed quantitative mass spectrometry. Nat. Protoc. 10, 1567–1593 (2015).

61. F. Weis, W. J. H. Hagen, Combining high throughput and high quality for cryo-electron microscopy data collection. Acta Crystallogr D Struct Biol. 76, 724–728 (2020).

62. D. N. Mastronarde, Automated electron microscope tomography using robust prediction of specimen movements. J. Struct. Biol. 152, 36–51 (2005).

63. A. Punjani, J. L. Rubinstein, D. J. Fleet, M. A. Brubaker, cryoSPARC: algorithms for rapid unsupervised cryo-EM structure determination. Nat. Methods. 14, 290–296 (2017).

64. T. Bepler, A. Morin, M. Rapp, J. Brasch, L. Shapiro, A. J. Noble, B. Berger, TOPAZ: A Positive-Unlabeled Convolutional Neural Network CryoEM Particle Picker that can Pick Any Size and Shape Particle. Microsc. Microanal. 25, 986–987 (2019).

65. A. Punjani, H. Zhang, D. J. Fleet, Non-uniform refinement: adaptive regularization improves single-particle cryo-EM reconstruction. Nat. Methods. 17, 1214–1221 (2020).

66. E. F. Pettersen, T. D. Goddard, C. C. Huang, E. C. Meng, G. S. Couch, T. I. Croll, J. H. Morris, T. E. Ferrin, UCSF ChimeraX: Structure visualization for researchers, educators, and developers. Protein Sci. 30, 70–82 (2021).

67. T. I. Croll, ISOLDE: a physically realistic environment for model building into low-resolution electron-density maps. Acta Crystallogr D Struct Biol. 74, 519–530 (2018).

68. A. Casañal, B. Lohkamp, P. Emsley, Current developments in Coot for macromolecular model building of Electron Cryo-microscopy and Crystallographic Data. Protein Sci. (2020) (available at https://onlinelibrary.wiley.com/doi/abs/10.1002/pro.3791).

69. S. R. Offley, M. M. Pfleiderer, A. Zucco, A. Fraudeau, S. A. Welsh, M. Razew, W. P. Galej, A. Gardini, A combinatorial approach to uncover an additional Integrator subunit. Cell Rep. 42, 112244 (2023).

70. P. V. Afonine, B. K. Poon, R. J. Read, O. V. Sobolev, T. C. Terwilliger, A. Urzhumtsev, P. D. Adams, Real-space refinement in PHENIX for cryo-EM and crystallography. Acta Crystallogr D Struct Biol. 74, 531–544 (2018).

71. J. Y. Young, J. D. Westbrook, Z. Feng, R. Sala, E. Peisach, T. J. Oldfield, S. Sen, A. Gutmanas, D. R. Armstrong, J. M. Berrisford, L. Chen, M. Chen, L. Di Costanzo, D. Dimitropoulos, G. Gao, S. Ghosh, S. Gore, V. Guranovic, P. M. S. Hendrickx, B. P. Hudson, R. Igarashi, Y. Ikegawa, N. Kobayashi, C. L. Lawson, Y. Liang, S. Mading, L. Mak, M. S. Mir, A. Mukhopadhyay, A. Patwardhan, I. Persikova, L. Rinaldi, E. Sanz-Garcia, M. R. Sekharan, C. Shao, G. J. Swaminathan, L. Tan, E. L. Ulrich, G. van Ginkel, R. Yamashita, H. Yang, M. A. Zhuravleva, M. Quesada, G. J. Kleywegt, H. M. Berman, J. L. Markley, H. Nakamura, S. Velankar, S. K. Burley, OneDep: Unified wwPDB System for Deposition, Biocuration, and Validation of Macromolecular Structures in the PDB Archive. Structure. 25, 536–545 (2017).

72. V. B. Chen, W. B. Arendall 3rd, J. J. Headd, D. A. Keedy, R. M. Immormino, G. J. Kapral, L. W. Murray, J. S. Richardson, D. C. Richardson, MolProbity: all-atom structure validation for macromolecular crystallography. Acta Crystallogr. D Biol. Crystallogr. 66, 12–21 (2010).

